# Identification of a Culturable Fungal Species and Endosymbiotic Bacteria in Saliva of *Aedes aegypti* and *Culex pipiens* and Their Impact on Arbovirus Infection *in Vitro*

**DOI:** 10.1101/2023.03.31.534949

**Authors:** Lanjiao Wang, Laure Remue, Nikki Adriaens, Alina Soto, Sam Verwimp, Joyce van Bree, Katrien Trappeniers, Leen Delang

## Abstract

Mosquito saliva plays a key role in arbovirus transmission and pathogenesis. This study isolated and identified culturable fungal and bacterial colonies from saliva harvested from *Aedes aegypti* (lab strain) and *Culex pipiens* (field-collected) mosquitoes. For the first time, *Penicillium crustosum* was identified in mosquito saliva. Culturable bacteria detected in mosquito saliva included *Serratia marcescens, Serratia nematodiphila*, *Enterobacter* spp., and *Klebsiella* spp., which were previously identified as mosquito or insect endosymbionts in the midgut or other organs. Analysis with 16S metagenomics showed that the bacterial community in saliva appeared more diverse than the bacterial communities in midguts. Blood feeding did not affect the fungal or bacterial load in mosquito saliva. Oral treatment of adult mosquitoes with antibiotics or an antifungal drug resulted in a significant reduction of resp. bacteria or fungi present in the mosquito saliva. Co-incubation of Semliki Forest virus with saliva from antibiotic or antifungal treated mosquitoes triggered a decrease in viral infection in human skin fibroblasts compared to non-treated saliva. This work lays the foundation for further exploration of the impact of fungi and bacteria in mosquito saliva on both vector competence and arbovirus infection in the mammalian host.

## Introduction

Mosquitoes of the *Culicidae* family are considered the deadliest animals on the planet (1), due to the ability of female mosquitoes to transmit pathogens such as arboviruses (*i.e.* arthropod-borne viruses) during blood feeding. The main mosquito vectors of medically important arboviruses, such as chikungunya virus (CHIKV), Zika virus (ZIKV), dengue virus (DENV), Yellow fever virus (YFV), and West Nile virus (WNV), are *Aedes aegypti* and *Culex pipiens* (1). Arboviruses are an emerging threat to the health of humans and animals worldwide (2). Symptoms caused by these viruses can vary from asymptomatic or mild febrile illness to severe diseases, such as chronic arthralgia and arthritis (CHIKV), congenital microcephaly and Guillain-Barré syndrome (ZIKV), dengue haemorrhagic fever and shock syndrome (DENV), and viral encephalitis (WNV), which can potentially be fatal (3–5). In recent decades, arbovirus epidemics have increased as a consequence of the development of global transportation systems, deforestation, urbanisation, and an increasing human population density.

Several decades of global efforts towards the development of arbovirus vaccines have resulted in few successful candidates. Furthermore, there are currently no approved, specific antiviral drugs available to treat arbovirus infections. Therefore, the current primary strategy to control arbovirus outbreaks in high-risk countries relies on mosquito vector control with insecticides aiming at reducing contact with potentially infected mosquitoes(6). Due to the long-term usage of insecticides of the same chemical classes, including organophosphates and synthetic pyrethroids, insecticide-resistant mosquitoes are widely spread and are considered a major threat to vector control by the World Health Organization (7). Other known limitations of using insecticides are ineffective timing of application, low efficacy, low community acceptance, and high economic costs (8). Therefore, new strategies for vector control are needed to reduce the global burden caused by arboviruses and other vector-borne pathogens. Promising results have been accomplished using host-associated microorganisms (*i.e.* microbiota members) as novel vector control strategies (9,10).

When feeding on an infected host, the female mosquito ingests an arbovirus simultaneously with the blood into the midgut. In order to be efficiently transmitted to another vertebrate host, the virus must replicate primarily in the midgut and then successfully disseminate to secondary organs, especially to the salivary glands and saliva. The virus must cross both the midgut and salivary gland barriers to reach and replicate in salivary glands. Finally, to infect a new host, the virus needs to be released into the mosquito saliva and deposited into the skin during a subsequent blood feeding (11). Hence, the transmission of viruses through mosquitoes among diverse hosts can be influenced not only by extrinsic factors such as temperature, humidity, and nutrition, but also by intrinsic mosquito-related factors including innate immune responses, microbiota, and genetic factors (12).

The midgut and the salivary glands have been characterised as key barriers to systemic arbovirus infection in the mosquito and to transmission within saliva. Meanwhile, both organs are also known to be colonised by dynamic microbial communities (13–16). Most studies have focused on how gut microbiota impact mosquito biology, including vector competence (17). Multiple interactions between microbial species and arboviruses have been demonstrated. For example, *Serratia odorifera*, a bacterium inhabiting the midgut of *Ae. aegypti*, has been shown to enhance mosquito susceptibility to DENV and CHIKV by suppressing the mosquito immune response (18,19). Furthermore, *Serratia marcescens* was identified as a midgut commensal bacterium with the ability to promote mosquito permissiveness to arboviruses by secreting SmEnhancin, a protein to digest the mosquito midgut membrane-bound mucins (20). In addition to bacteria, a *Talaromyces* fungus was found in the midgut of field-captured *Ae. aegypti*, rendering the mosquito more permissive to DENV infection by down-regulating digestive enzyme genes and their activity in the midgut (21). Although the gut forms a first important barrier in establishing a viral infection in the mosquito, viral transmission to the next mammalian host occurs via the mosquito saliva. While biting a host, female mosquitoes probe into the host skin in search of a blood vessel and meanwhile inject saliva into the epidermis and dermis of the vertebrate host. Mosquito saliva contains vasodilatory, anti-hemostatic, angiogenic, and anti-inflammatory molecules that ensure continuous blood pumping to the mosquito during feeding (22,23). Additionally, mosquito saliva contains molecules that could alert the immune system of the vertebrate host and potentially block viral transmission. Therefore, mosquito saliva components are being studied as promising candidates for vaccine development (24). Interestingly, two recent papers demonstrated that the saliva of *Aedes albopictus* and *Anopheles* mosquitoes contains bacteria (25,26). Nevertheless, our current understanding of which microbiota members are present in the saliva of *Aedes* and *Culex* mosquitoes is still very limited. Furthermore, it is unclear whether and how the saliva microbiota impacts vector competence and, being transferred to the mammalian host, if they downstream affect the virus replication and virus infection in the host.

Here, we describe that both fungi and bacteria can be cultured from the saliva of laboratory-maintained *Ae. aegypti* and field-collected *Cx. pipiens* mosquitoes. In addition, oral treatment of adult mosquitoes with antibiotics or an antifungal drug resulted in a significant reduction of bacteria or fungi in saliva. Finally, we demonstrated that saliva from antibiotic or antifungal treated mosquitoes decreased infectious viral loads of Semliki Forest virus (SFV) in human skin fibroblasts Further *in vivo* studies are required to better understand its impact on viral infection.

## Results

### Identification of fungi in mosquito saliva

Mosquito saliva pooled from 8 to 12 *Ae. aegypti* mosquitoes was cultured on LB agar plates for at least 5 days at 28°C. Fungi appeared as grey-blue powdery or fuzzy colonies (indicated with black arrows, **Fig 1A**), distinguishable from bacterial colonies. The fungi colony forming units (CFU) per mosquito were recorded after culturing for at least 120h. No fungal growth was observed in the negative environmental control, which contained the blank salivation solution present on the bench during collecting the saliva. Next, saliva collected from single *Ae. aegypti* and field-collected *Cx. pipiens* female mosquitoes were cultured in similar conditions. 40% of *Ae. aegypti* (12/30 females) and 33% of *Cx. pipiens* (7/21 females) had culturable fungi in their saliva (**Fig 1B**). ITS3-4 universal qPCR, which determined the fungal load, was performed to verify the presence of fungal DNA in saliva, salivary glands, and midgut samples of *Ae. aegypti*. The 17S mosquito housekeeping gene was used as an internal control. The highest fungal load was detected in midgut samples, with 4.5×10^6^ median ITS copies/mosquito (**Fig 1C**). Salivary glands contained a fungal load of 3.0×10^3^ median ITS copies/mosquito, higher than the load in saliva samples (83 median ITS copies/mosquito). The latter was not significantly different from the fungal load in the negative control.

**Figure 1.**
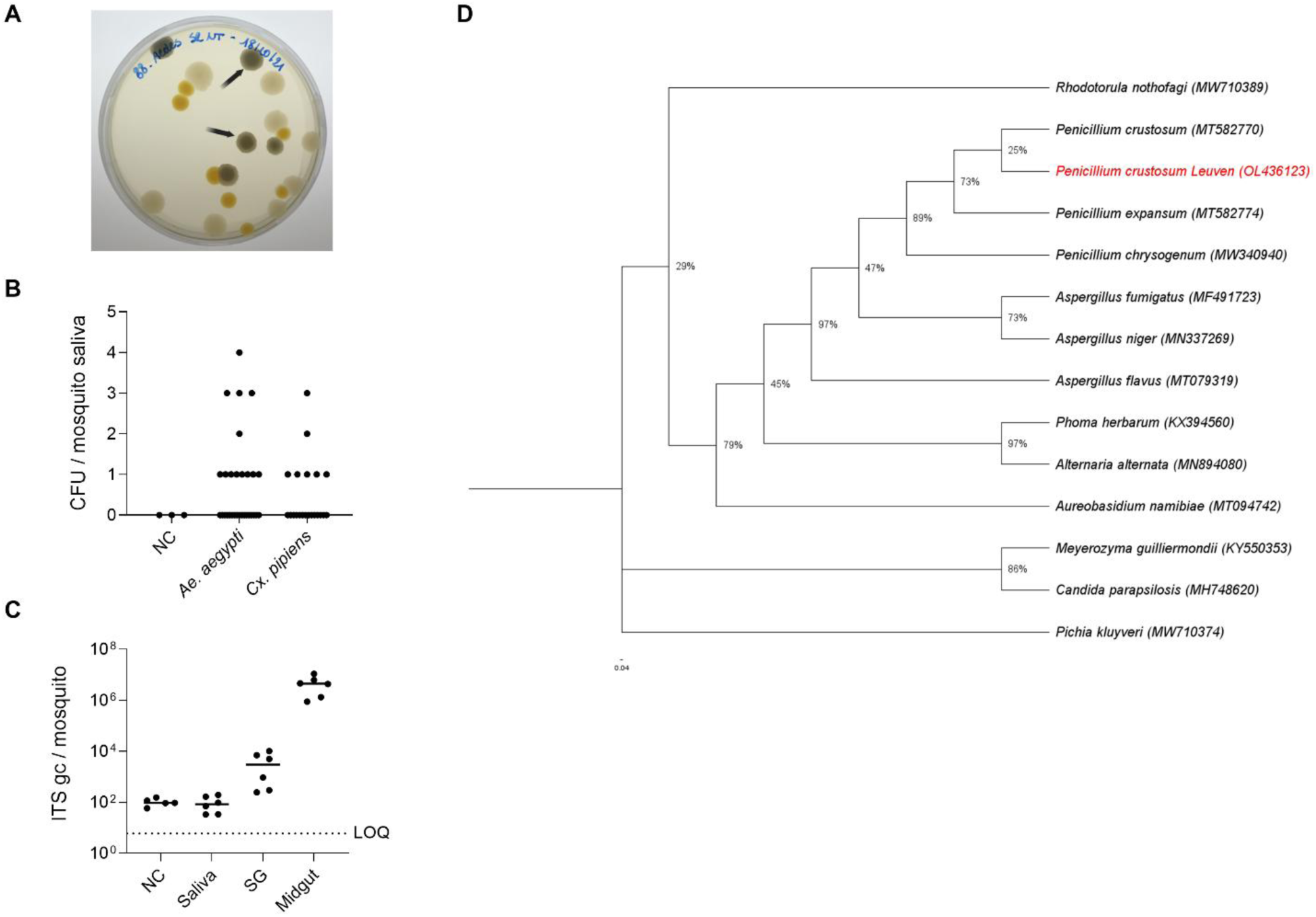
Presence of fungi in the saliva of *Aedes aegypti* and *Culex pipiens* mosquitoes. **(A)** Mosquito saliva of *Ae. aegypti* (lab strain) was harvested by forced salivation and cultured on LB agar plates for 120 h at 28°C. Fungal colonies are representatively indicated by black arrows. **(B)** Mosquito saliva from single *Ae. aegypti* (lab strain) and *Cx. pipiens* (field) mosquitoes were cultured on LB agar plates. The number of CFU/ mosquito was quantified (by visual counting) after culturing for 120 h at 28°C. Each dot represents one pool of 8-12 saliva samples normalized per mosquito. Lines show the median value of the tested samples. **(C)** Genome copies of fungi in different organs of *Ae. aegypti* were quantified by ITS qPCR. Negative control (NC) consisted of the blank solution used as environment control for forced salivation. Each point represents an independent pool containing samples harvested from 8-12 mosquitoes normalized per mosquito. Lines show the median value of the tested samples. LOQ presents the limit of quantification of the qPCR. SG= salivary glands. **(D)** Maximum likelihood tree was constructed using MEGA-X with the substitution model Kamura 2-parameter model (K2) and Gamma distributed (G) (500 bootstrap replicates), by using the ITS3-4 segments of *Penicillium crustosum* identified in mosquito saliva (OL436123; isolated from *Ae. aegypti* and *Cx. pipiens*) and other fungal species from GenBank which were previously detected in mosquito body and organs.

To identify the fungal colonies cultured from *Cx. pipiens* or *Ae. aegypti* saliva on agar plates, DNA was extracted from individual colonies and subjected to a universal ITS3-4 fungal identification PCR (27) and subsequent Sanger sequencing analysis. The results showed a 99% identity similarity by BLAST (NCBI) for both *Penicillium expansum* and *Penicillium crustosum*. Therefore, a specific PCR for *Penicillium crustosum* was performed(28). An 892 bp segment could be amplified by *P. crustosum*-specific primers from all samples (**Fig S1**), indicating that the fungal colonies were *Penicillium crustosum* rather than *Penicillium expansum*.

A phylogenetic analysis was performed to compare the ITS3-4 segments of *Penicillium crustosum* detected in the mosquito saliva from this study to the *Penicillium crustosum* and *Penicillium expansum* species previously reported in GenBank (**Fig 1D**). The bootstrap values indicate how many times (out of 500 repeats) the sequences clustered together when repeating the tree construction on a re-sampled data set. In 89% of the possibilities, the *Penicillium* clade, including the species identified in the mosquito saliva, was related to the previously described mosquito inhabitant *Penicillium chrysogenum* (29). In 97% of the possibilities, the *Penicillium* clade was related to a clade containing *Aspergillus fumigatus, Aspergillus niger* and *Aspergillus flavus*, which were previously reported in mosquitoes (30).

### Identification of bacteria in mosquito saliva

Culturing of mosquito saliva bacteria was performed on LB agar plates, as no significant difference was obtained when comparing the bacterial load and diversity of mosquito saliva bacteria using LB agar plates or BHI/blood agar plates (**Fig S2, Table S1 and S2**). For the culturing, saliva from 8-12 mosquitoes was pooled and 50 µl of the pool was inoculated on agar plates. In a first step, bacteria were characterized by morphological features (*e.g.* shape and colour) (**Fig 2A**, bacteria indicated with black arrows). The bacterial CFU/mosquito was defined at 48h post-culture at 28°C for both *Ae. aegypti* and *Cx. pipiens* (**Fig 2B**). No colonies grew in the environmental control, containing only the salivation solution placed on the bench during the salivation procedure. To further support this, ‘salivation’ was also performed using dead *Aedes aegypti* females (previously killed at –20 °C for 5 minutes) or by cutting the proboscis of live females and immersing it in 20 µL of blank solution, the collected saliva was cultured under the same conditions, and no bacterial or fungal colonies were observed. Furthermore, no colonies were observed on the plates with salivary gland samples from the same mosquitoes (**Fig S3**). Using 16S qPCR, the highest bacterial load was found in the midgut of *Ae. aegypti* (10^4^ median copies/mosquito), followed by saliva (170 median copies/mosquito) and salivary glands (38 median copies/mosquito) (**Fig 2C**). Individual bacterial colonies were picked from the plates based on their characteristics (shape, colour and size) and were identified by 16S Sanger sequencing (**Fig 2D**). *Klebsiella* spp.*, Enterobacter* spp.*, Asaia* spp.*, Serratia marcescens*, *Serratia nematodiphila, Microbacterium* spp., and *Chryseobacterium* sp. were identified in the saliva of *Ae. aegypti* (indicated with ○ in **Fig 2E** and listed in **Table S1**), whereas only *Serratia marcescens* and *Serratia nematodiphila* were found in the saliva of *Cx. pipiens* mosquitoes (indicated with Δ in **Fig 2E** and listed in **Table S2**). Phylogenetic analysis showed that, in 98% of the possibilities, *Serratia* sp. sequences of this study clustered with *Serratia* sp. (FJ372764) previously found in mosquitoes (**Fig 2E**) (31). Additionally, the clade containing *Enterobacter* spp. And *Klebsiella* spp. was related to those previously identified in the soil where the mosquitoes resided (OK035565 and OL441036) in 87% and 89% of the possibilities, respectively. In 74% of the possibilities, the *Asaia* spp. detected in saliva were closely related to the *Asaia* sp. that was previously detected in mosquitoes (FJ372761) (31). Finally, the *Chryseobacterium* sp. and *Microbacterium* sp. were found in 100% of the possibilities related to the species previously observed in mosquitoes (KY002136 and OM033360).

**Figure 2.**
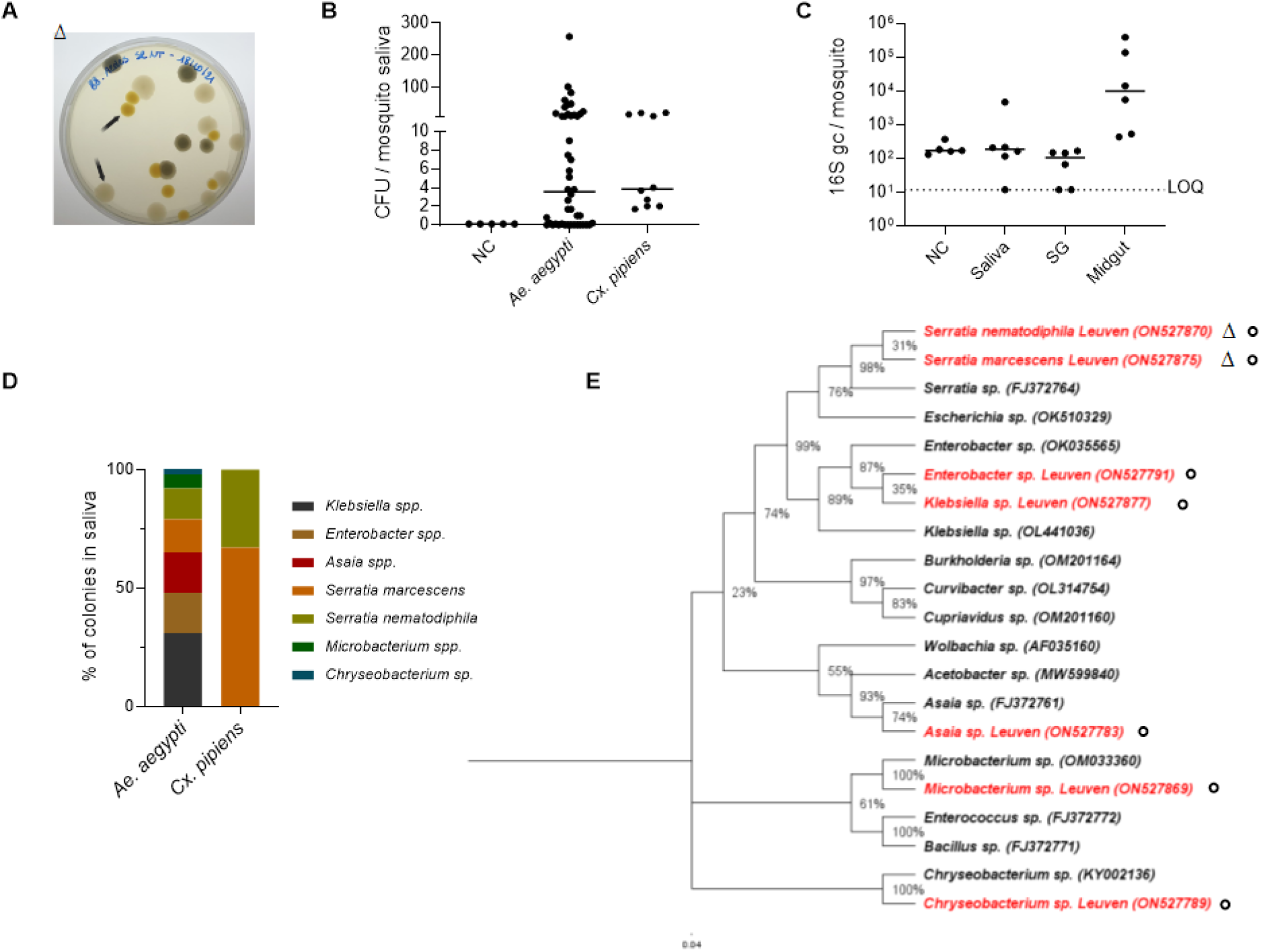
Presence of bacteria in the saliva of *Aedes aegypti* and *Culex pipiens* mosquitoes. **(A)** Mosquito saliva of *Ae. aegypti* (lab strain) was harvested by forced salivation and cultured on LB agar plates. Bacterial colonies are representatively indicated by black arrows. **(B)** Mosquito saliva pools from both *Ae. aegypti* (lab strain) and *Cx. pipiens* (field collection) were cultured on LB agar plates at 28°C. The CFU/saliva pool was quantified (by visual counting) after culturing for 48 h. Each dot represents one pool of 8-12 saliva samples normalized per mosquito. Lines show the median value of the tested samples. **(C)** Genome copies of the 16S bacterial gene in different organs of *Ae. aegypti* were quantified by 16S-qPCR. NC as environmental control consisted of the solution used for forced salivation. Each point represents an independent pool containing samples harvested from 8-12 mosquitoes normalized per mosquito. Lines show the median value of the tested samples. LOQ presents the limit of quantification of the qPCR. SG= salivary glands. **(D)** Identification by 16S Sanger sequencing of the individual bacterial colonies (based on shape, colour, and size) detected on the agar plates after cultivating *Ae. aegypti* and *Cx. pipiens* saliva. **(E)** ML tree was constructed using MEGA-X with the substitution model being Kamura 2-parameter model (K2) and Gamma distributed (G) (500 bootstrap replicates), by using the Sanger sequences of bacterial colonies identified in mosquito saliva (*Ae. aegypti* indicated with ○ and *Cx. pipiens* indicated with Δ) and other species from GenBank which were previously detected in mosquitoes.

To profile the bacterial composition, 16S metagenomics analysis was performed on saliva, salivary gland, and midgut sample pools of *Ae. aegypti* females without a prior cultivation step. DNA extraction of the saliva samples resulted in a relatively low DNA yield (around 10 ng/µL). Approximately 10,000 reads were obtained per saliva pool, with a total read base of 70 Mbp/sample pool and Q30 of more than 82%. The sequencing outputs of saliva samples (1.1×10^5^ reads, SD±1.24×10^4^) were equal or slightly higher than the ones of midguts (9.5×10^4^ reads, SD±1.3×10^4^) and salivary glands (1.1×10^5^ reads, SD±4.1×10^3^) (**Table S3**). There was no significant systematic difference in alpha-diversity among saliva, midguts, and salivary gland samples (Kruskal-Wallis chi-squared = 3.5043, df = 3, p-value = 0.3202) (**Fig 3A**). However, ordination picked out a clear separation between the saliva samples and the other organ samples (**Fig 3B**). The taxonomic distribution of the top 15 bacterial sequences of saliva, salivary glands, and midguts was generated at the genus level (**Fig 3C**). Two dominant genera in salivary glands were *Pseudomonas* (55% ± 27%) and *Cutibacterium* (5% ± 3%). In saliva samples, genera appeared more diversely, with the most relatively dominant genera being *Delftia* (21% ± 18%), *Methylobacterium*-Methylorubrum (5% ± 3%), *Flavobacterium* (7%±5%) and *Chryseobacterium* (6%±5%). In midguts, the top three genera were *Enterobacter* (36% ± 14%), *Pseudomonas* (13% ± 4%) and *Cutibacterium* (11% ± 7%). Interestingly, the most dominant genus in the blank control was *Enhydrobacter* (19%), which was rarely found in the mosquito samples.

**Figure 3.**
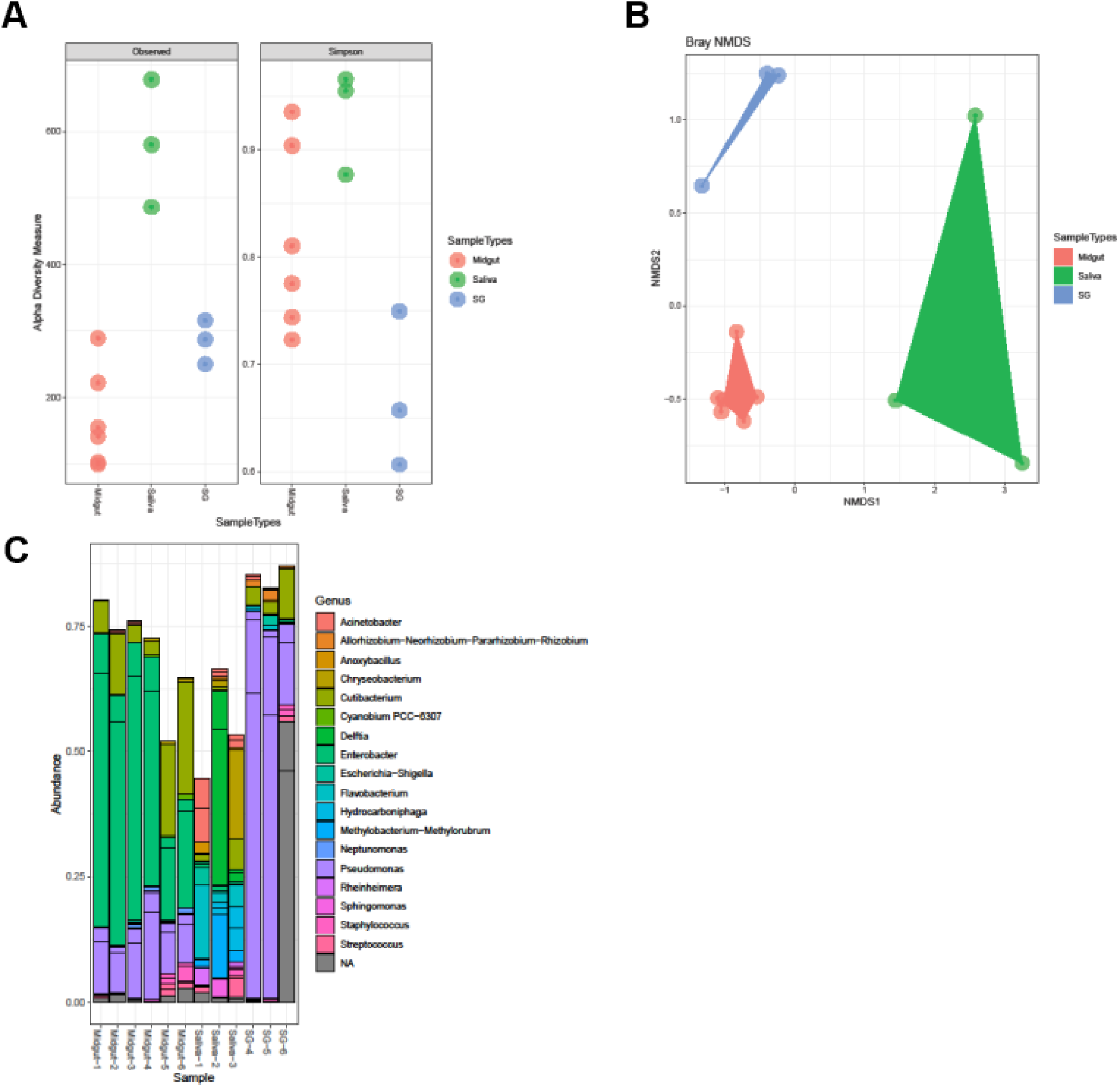
Bacterial richness and diversity in saliva, midguts and salivary glands of *Aedes aegypti* mosquitoes. The 16S rRNA V3-V4 metagenomic sequencing was analyzed in DADA2, **(A)** alpha diversity in the microbial communities identified from different organs was visualized by Phyloseq and scored using the “observed” and “Simpson” diversity indexes; **(B)** Ordination was assessed using Bray-Curtis distances; **(C)** Taxonomic distribution of the top 20 sequences was shown at the genus level, SG = salivary glands.

### Effect of blood feeding, life stages, and sex on the mosquito microbiota

To assess whether blood feeding could influence the presence of the saliva microbiota, culturable fungi and bacteria were determined in the saliva of *Ae. aegypti* following a blood meal. Our results showed that the bacterial and fungal loads were not different in the saliva of fed versus non-fed mosquitoes (**Fig 4A-B**), demonstrating that a blood meal did not alter the microbiota loads in the saliva. Of note, no fungal or bacterial colonies were observed after one week of culture of the blood itself, indicating there were no input fungi or bacteria from the bloodmeal.

**Figure 4.**
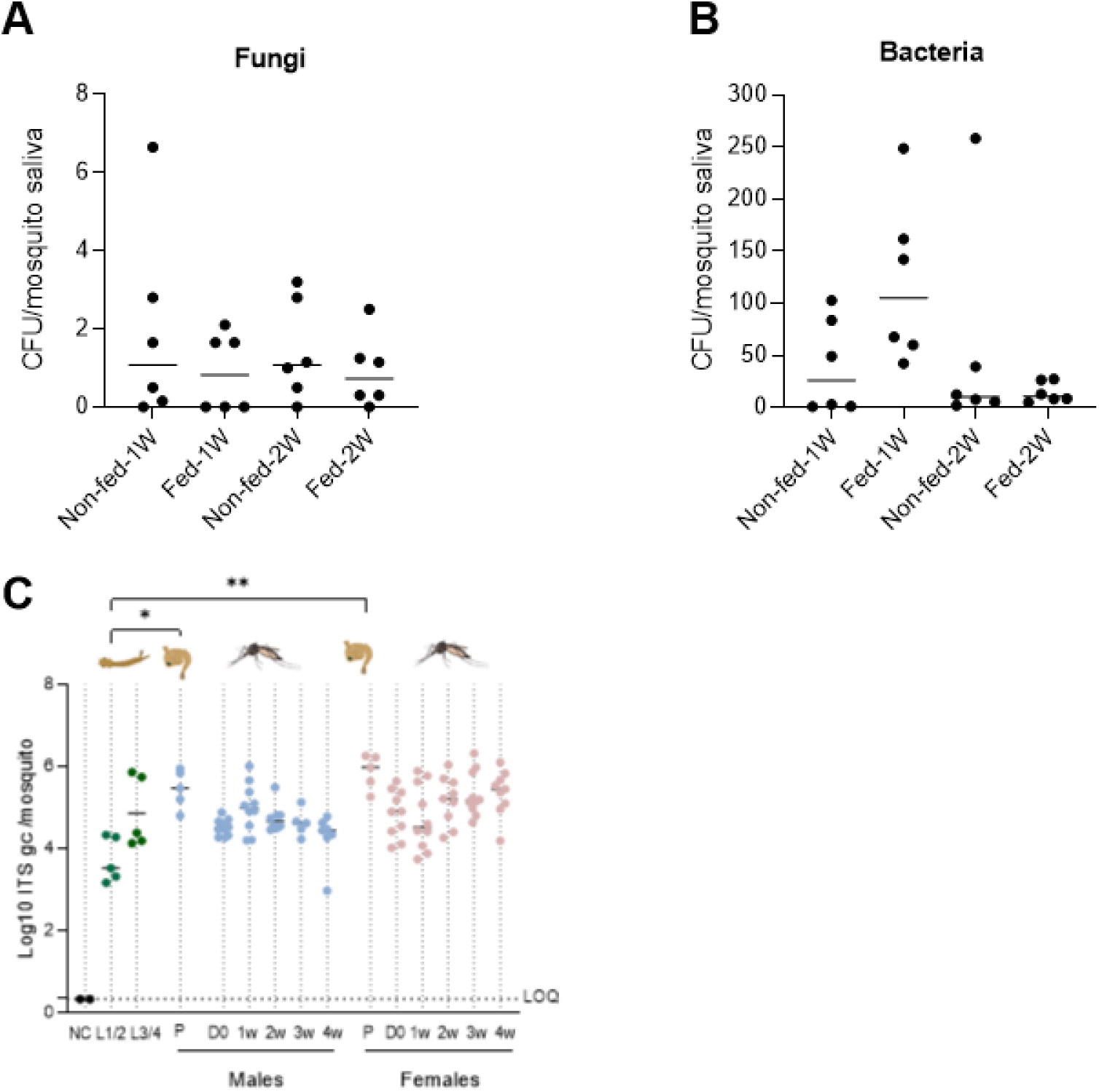
The effect of blood feeding and life stages on the microbiota. **(A)** Bacterial and **(B)** fungal CFU of saliva from one-week-old female *Ae. aegypti* adults fed on rabbit blood. Saliva was harvested after one week and two weeks post-blood meal. The CFU/saliva pool was quantified (by visual counting) after culturing. Each dot represents one pool of 8-12 saliva samples normalized per mosquito. Lines show the median value of the tested samples. The loads were compared between fed and unfed groups. Statistical analysis was performed using the Mann-Whitney test. **(C)** Genome copies of fungi at different life stages of *Ae. aegypti* were quantified by ITS qPCR using homogenized mosquito bodies. The lines represent the median values. LOQ presents the limit of quantification of the qPCR. D0: newly emerged adults, Negative control (NC) consisted of the blank solution used as environment control for forced salivation. Statistical analysis was performed using the Kruskal-Wallis test (*, p<0.05; **, p<0.01).

The effect of the mosquito’s life stages on its microbiota in *Ae. aegypti* was determined with qPCR in larvae, pupae and female/male adults (**Fig 4C** **and FigS4**). The fungal loads increased during the immature life stages from 3.3×10^3^ median ITS copies/mosquito in L1/L2 larvae to 3.0×10^5^ median ITS copies/mosquito in male pupae (Kruskal-Wallis test; p<0.05) and to 1.0×10^6^ median ITS copies/mosquito in female pupae (Kruskal-Wallis test; p<0.01) (**Fig 4C**). A drop in the fungal load was observed during the metamorphosis of male and female pupae into newly emerged male and female mosquitoes, although non-significant (Kruskal-Wallis test; ns, *p*>0.05). There was no significant difference in fungal loads between adult males and females (Kruskal-Wallis test; ns, *p*>0.05).

In contrast to fungi, the bacterial loads in immature life stages were already high and did not increase further in the adult life stage. Indeed, no significant difference was observed between larvae or pupae compared to adult mosquitoes for the bacterial loads (**Fig S4**). A similar drop was observed during metamorphosis in bacterial genome copies from pupae to newly emerged adult mosquitoes, although non-significant (Kruskal-Wallis test; ns, *p*>0.05). The bacterial load in female mosquitoes was higher compared to the loads in male mosquitoes, with a statistically significant difference in median bacterial colonies of 6.5×10^4^ CFU/mosquito between 2-weeks old males and females (Kruskal-Wallis test; *p*<0.01). At the end of their life, 4-weeks old females also had a significantly higher median CFU/mosquito of 1.8×10^5^ compared to 4-weeks old males (1.5×10^3^ CFU/mosquito) (Kruskal-Wallis test; *p*=0.04) (**Fig S4**). **Supplementary Figure S5** shows the culturable bacteria identified by Sanger sequencing in *Ae. aegypti* at different life stages in males and females.

A correlation analysis was performed for male and female mosquitoes to investigate the relationship between bacterial and fungal loads for the adult life stages. Interestingly, an opposite correlation was observed in male compared to female mosquitoes. In male mosquitoes, a significant negative correlation was observed between bacterial and fungal loads (Pearson Correlation; R2=0.3281; p<0.05). This showed that male mosquitoes with a high bacterial load had lower fungal loads (**Fig 5A**). This was in contrast to the significant positive correlation in females between bacterial and fungal loads (Pearson Correlation; R2=0.1395; p<0.05) (**Fig 5B**). Female mosquitoes with a high bacterial load, also had a high fungal load. The difference in correlation between male and female could be linked to the blood feeding based physiology of females, although this requires further investigation.

**Figure 5.**
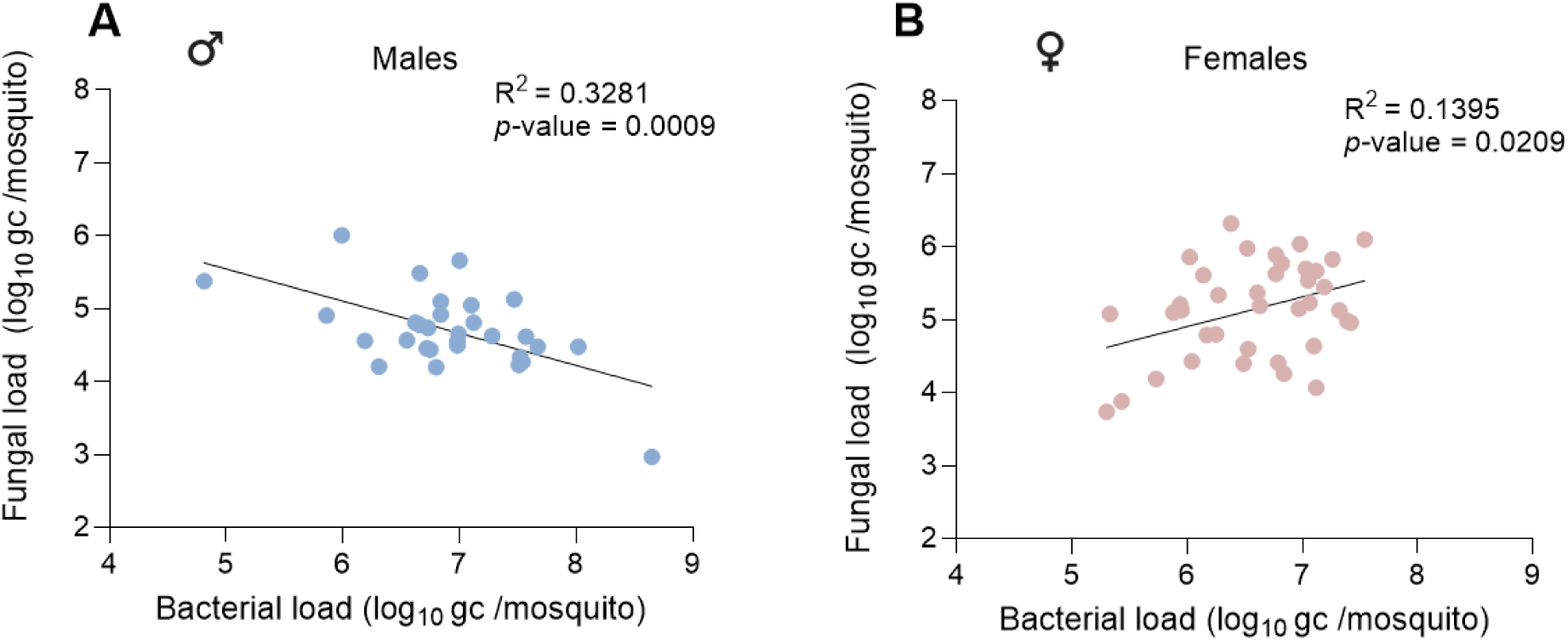
Correlation between fungal and bacterial loads in mosquito (A) males and (B) females. Genome copies of the 16S bacterial gene of *Ae. aegypti* homogenized mosquito bodies were quantified by 16S-qPCR. Genome copies of fungi in of *Ae. aegypti* homogenized mosquito bodies were quantified by ITS qPCR. Each point represents an independent pool containing samples harvested from 8-12 mosquitoes normalized per mosquito. Statistical analysis was performed using the Pearson Correlation test.

### Depletion of culturable fungi and bacteria in mosquito saliva after oral treatment of adult mosquitoes

In order to obtain mosquito saliva devoid of fungi or bacteria, adult mosquitoes were given respectively an oral antifungal or antibiotic treatment. To deplete fungi, female *Ae. aegypti* of 5-7 days old were provided with 10% sterile sugar solution supplemented with 25 µg/mL Fungin for 10 days (**Fig 6A**). Antifungal treatment of the adult mosquitoes resulted in the complete absence of fungi in the saliva, whereas the bacterial load remained unaffected (**Fig 6C**). To deplete bacteria, females were provided with 10% sterile sugar solution supplemented with 20 units/mL of penicillin and 20 µg/mL streptomycin for 6 days, as described previously (20). Oral antibiotic treatment significantly reduced the bacterial load in the saliva (Mann-Whitney test, p=0.028), but not in a complete depletion (**Fig 6B**). Remaining bacteria following this treatment were identified as *C. cucumeris, M. maritypicum*, *M. oxydans* and *S. maltophilia*, representing 17%, 69%, 12% and 2.7% of all colonies, respectively. Therefore, gentamycin was added to the antibiotic cocktail, a drug effective against both gram-positive and gram-negative bacteria. The addition of gentamycin resulted in full depletion of culturable bacteria in the saliva (**Fig 6B**). Different doses of penicillin, streptomycin, and gentamycin were evaluated to optimise the treatment. In addition, oral antifungal and antibiotic treatments at different doses were shown not to affect the survival of females (**Fig S6**). As it was possible to obtain saliva devoid of bacteria or fungi after orally treating the mosquitoes, we concluded that the culturable bacteria and fungi detected in saliva originated from the mosquito body itself and not from the environment or the mosquito outer body.

**Figure 6.**
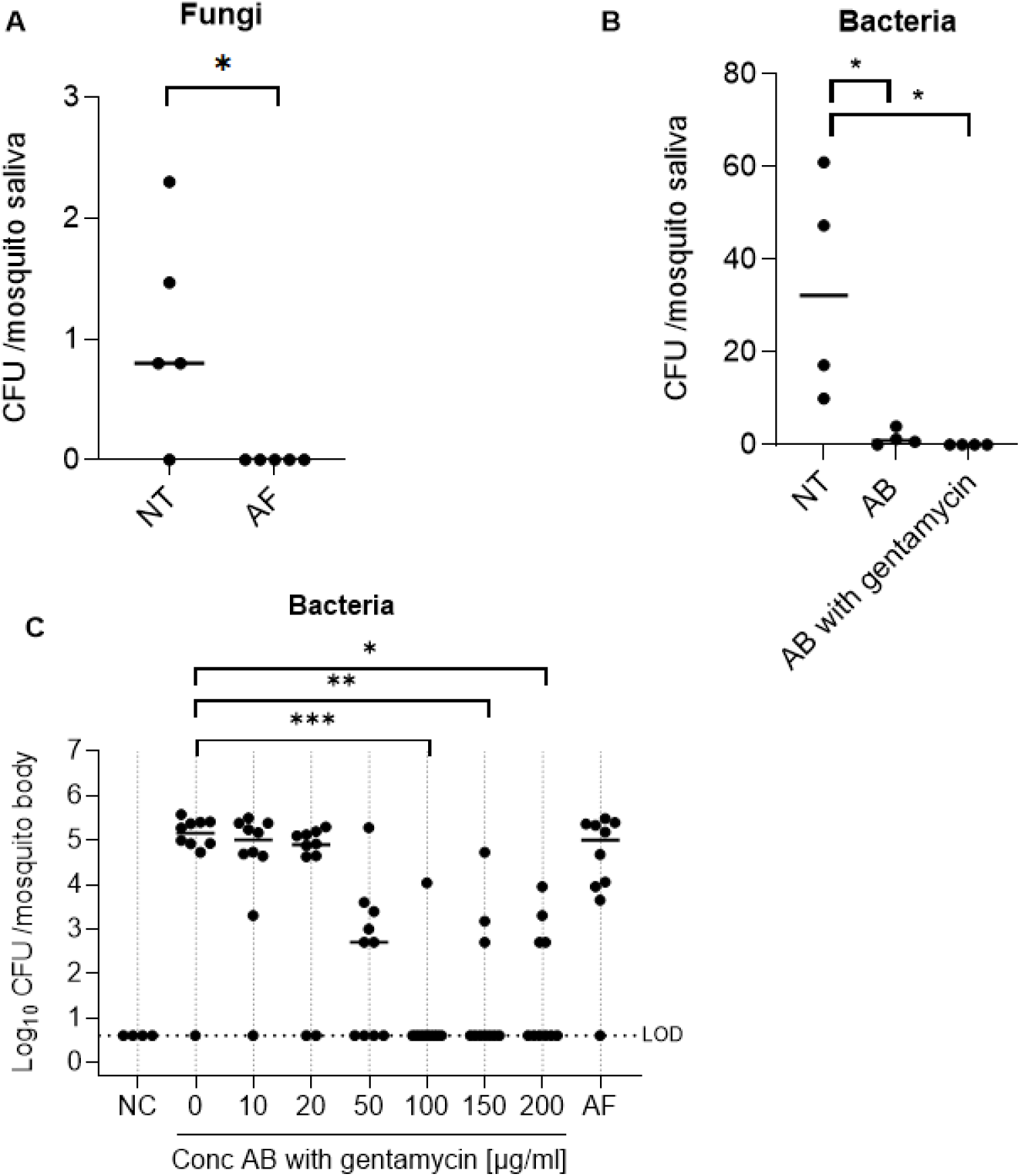
Depletion of culturable fungi or bacteria in mosquito saliva after oral treatment. **(A)** Fungal load in saliva from antifungal-treated mosquitoes (AF; 25 µg/mL of Fungin in 10% sucrose for 10 days) and non-treated saliva (NT). The CFU/saliva pool was quantified (by visual counting) after culturing. Each dot represents one pool of 8-12 saliva samples normalized per mosquito. **(B)** Bacterial load in saliva from antibiotics-treated mosquitoes (AB; 20 units/mL of penicillin and 20 µg/mL streptomycin in 10% sucrose for 6 days with and without 100 µg/mL gentamycin) and non-treated saliva (NT). The CFU/saliva pool was quantified (by visual counting) after culturing. Each dot represents one pool of 8-12 saliva samples normalized per mosquito. **(C)** Female *Ae. aegypti* adults (5-7 days old) were divided between three treatment groups: untreated control group (NT: only 10% sucrose), antifungal-treated (AF: 25 µg/mL of Fungin in 10% sucrose for 10 days) and antibiotic-treated (AB: serial dilution up to 200 units/mL of penicillin and 200 µg/mL streptomycin + gentamycin in 10% sucrose for 6 days). Bacterial loads in the individual bodies were quantified by determining CFU/mosquito. LOD presents the limit of detection. Significant differences were demonstrated by the Mann-Whitney test (two groups) or Kruskal-Wallis test (≥ three groups) (*, *p*<0.05; **, *p*<0.01; ***, *p*<0.001).

### Reduced viral titers after co-incubation of Semliki Forest virus with saliva of antibiotic or antifungal treated mosquitoes

To investigate the potential impact of fungi and bacteria in the mosquito saliva on arbovirus replication *in vitro*, saliva was collected from antifungal-treated females (AF) or antibiotic-treated females (AB) and from non-treated female *Ae. aegypti* (NT) (**Fig 7A-C**). Additionally, filtered saliva, from which fungi and bacteria were removed physically using a 0.2 µm filter, was tested. Interestingly, the direct inoculation of human skin fibroblasts with SFV and AB saliva or AF saliva significantly reduced SFV replication compared to the NT saliva (Kruskal-Wallis test; *p*<0.01) (**Fig 7A**). This inhibition was not observed when skin fibroblasts were incubated with filtered saliva (**Fig 7A**). Similarly, pre-incubation of saliva and SFV for 1h prior to inoculation on human skin fibroblasts, significantly lowered SFV replication for AB saliva or AF saliva compared to NT saliva (Kruskal-Wallis test; *p*<0.01). Again, there was no difference between NT saliva and filtered saliva (**Fig 7B**). Furthermore, in both conditions, no difference was observed between SFV infectivity in the presence or absence of NT saliva. Finally, saliva was pre-incubated with skin fibroblast cells for 1h, after which the cells were infected with SFV. No significant difference was observed between the NT saliva, AF saliva or AB saliva conditions; however, SFV replication was significantly reduced when cells were pre-incubated with filtered saliva (Kruskal-Wallis test; *p*<0.01) compared with NT saliva (**Fig 7C**).

**Figure 7.**
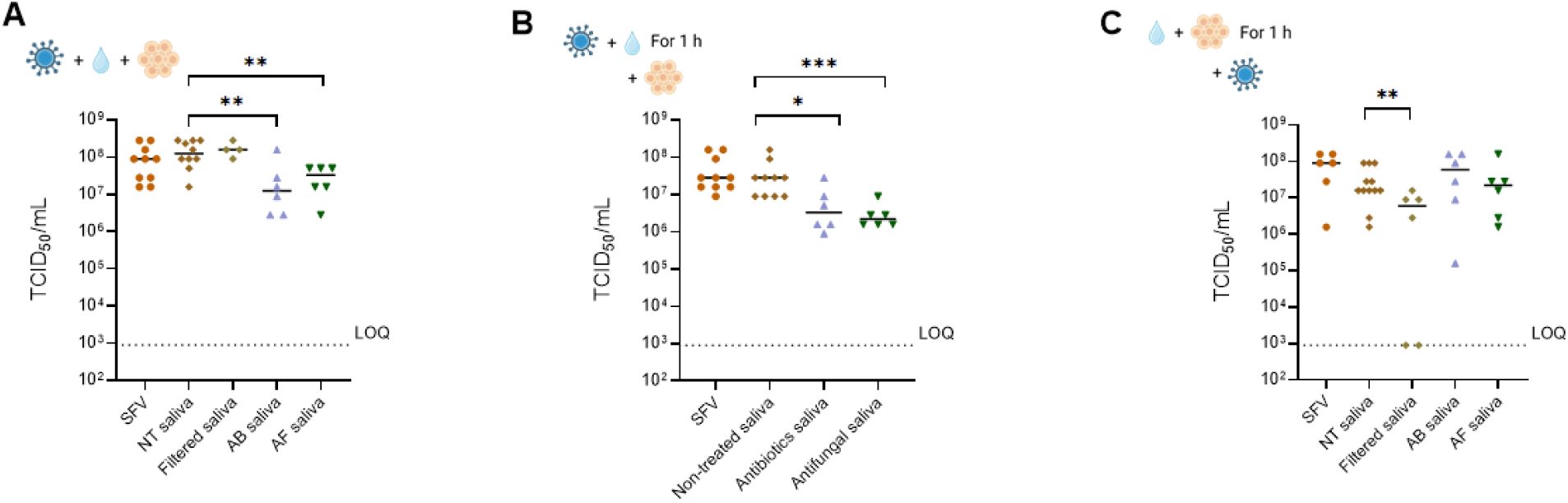
Effect of mosquito saliva on Semliki Forest virus infectivity in human skin fibroblasts. SFV infectious titers following exposure to mosquito saliva. For the positive control (SFV group), saliva was replaced by 1X PBS. **(A)** Human skin fibroblasts were directly inoculated with saliva (saliva from 5 females/well) and SFV (MOI:0.001), **(B)** SFV (MOI:0.001) was pre-incubated with saliva for 1 h at 28°C before inoculation on human skin fibroblasts, **(C)** Skin fibroblast cells were preincubated with saliva 1 h before infection with Semliki Forest virus (MOI: 0.001) and cells were washed three times. The cell supernatant was harvested at 24 h post-infection and the viral yield was determined by end-point titrations on Vero cells. TCID_50_/ml was calculated using the Reed and Muench method. Kruskal-Wallis test was used for statistical analysis (ns, p>0.05; *, p<0.05; **, p<0.01; ***, p<0.001). The dotted line indicates the limit of quantification (LOQ).

## Discussion

Mosquito saliva is a complex mixture of protein, lipid, and microbial components; however, the latter has not been studied in great detail. Here, we aimed to explore the presence of bacteria and fungi in *Aedes* and *Culex* mosquito saliva. Most saliva bacteria identified in this study were previously reported in either the midgut or other mosquito organs (**Fig 2**). Identified gram-negative genera were *Asaia*, *Enterobacter*, *Klebsiella*, *Chryseobacterium,* and *Serratia*. Interestingly, *Asaia*, belonging to the *Acetobacteraceae* family, was previously shown to colonize the salivary glands and the midgut of mosquito species such as *Aedes aegypti, Aedes albopictus, Anopheles stephensi,* and *Anopheles gambiae* (13,32). Importantly, *Asaia* has been considered a potential vector of anti-plasmodial factors through a paratransgenic approach, where the symbiotic bacterium acts as a Trojan horse to produce components that target the pathogen(33). The bacterial genera *Enterobacter* and *Klebsiella*, belonging to the family *Enterobacteriaceae,* are both frequently found in the gut of adult *Aedes* spp. and *Cx. Pipiens* (16,34). Furthermore, *Serratia* of the *Yersiniaceae* family was identified in the salivary glands and the midgut of *Aedes* and *Anopheles* mosquitoes (13) and the midgut of *Culex* mosquitoes(34). Interestingly, *Serratia marcescens* was found to have the ability to render mosquitoes more permissive to arbovirus infection (20). *Chryseobacterium,* a member of the *Weeksellaceae* family, was detected in all life stages of *Ae. aegypti* (35), which is in line with our observations that this species was found in both pupae and saliva samples of adult females. *Microbacterium,* belonging to the family *Microbacteriaceae,* was reported to be abundantly present in the larval stage of *Ae. Aegypti* (36). In this study, these gram-positive bacteria (for example *M. maritypicum* and *M. oxydans*) were mainly identified in the pupal stage, but also in *Ae. aegypti* adults, suggesting that a set of bacteria such as *Chryseobacterium* and *Microbacterium* might be conserved in *Ae. aegypti* colonies, despite being maintained in different laboratories (37).

Our results on the saliva microbiota are in line with previous findings showing the presence of bacteria in saliva from *Ae. albopictus* and *Anopheles* mosquitoes (25,26). Intriguingly, the same bacterial genera were identified in the saliva of *Ae. aegypti* (Paea strain) reared in our laboratory as in the saliva of *Anopheles gambiae* (G3 strain, MR4, MRA-112) and *Anopheles stephensi* (Sind-Kasur Nijmegen strain) reared in a laboratory in Italy (38). The 16S NGS data obtained in this study showed that the bacterial community in saliva samples was more diverse than in midgut samples. These findings are consistent with previous research (39), which showed that *Ae. albopictus* saliva harboured a richer microbial community compared to its midguts.

Our study does not allow to distinguish the effect of mosquito species (*Aedes* vs *Culex* mosquitoes) from their adaptation to the lab (field collected vs lab-reared mosquitoes) as we only used *Aedes* lab mosquitoes and *Culex* field mosquitoes. As a follow-up, an in-depth comparison between field and lab mosquitoes of the same species (*i.e.* mosquitoes reared for an extended period under standard conditions versus those recently transferred from the field to the lab), as well as further examination of the effects related to mosquito species would be of interest. Notably, Gómez *et al.* 2025 very recently compared the bacterial species in lab-reared and field-collected *Ae. aegypti* in Colombian populations, showing that *Pseudomonas* and *Acinetobacter* were present in both conditions and that field mosquitoes harbored a more diverse microbiota compared to lab mosquitoes (40).

In *Aedes* and *Culex* mosquitoes, fungal communities are mainly composed of Ascomycota (73-92%) and Basidiomycota (8–25%), including yeasts and filamentous species (41). Several species of filamentous fungi (*e.g. Aspergillus gracilis*, *Penicillium sp.*) and yeasts (*e.g. Candida sp*.) were highly detected. These fungi were shown to play a role in the development, survival and reproduction of mosquitoes (42). Previously, *Penicillium* spp. were identified in other mosquito species or habitats such as the breeding water (43). For instance, in Iran, *Penicillium* spp. were found to be the most abundant entomopathogenic fungi isolated and identified from larvae and breeding water of *Anopheles* and *Culex* mosquitoes (44). In Brazil, *Penicillium citrinum* was identified in *Aedes* larvae obtained from natural and artificial breeding sites (45). *Penicillium chrysogenum* was found in all life stages of *Cx. pipiens* in Egypt (46) and is particularly of interest as *P. chrysogenum* isolated from the midgut of field *Anopheles* mosquitoes rendered the mosquito more susceptible to a *Plasmodium* infection via suppressing the mosquito’s innate immune system (47). Here, we isolated and identified for the first time the fungus *Penicillium crustosum* from the saliva of both an *Ae. aegypti* lab strain and *Cx. pipiens* field-collected mosquitoes (**Fig 1**). *P. crustosum* is a common blue mold, morphologically characterized by typically gray-blue-green conidia. This species has been frequently detected in fruits (*e.g.* peaches, apples, pears), wheat, peanuts and other nuts, propolis, and food waste (48,49). It is a food–borne ubiquitous fungal species which produces roquefortine C and penitrems (50). Interestingly, penitrems have demonstrated insecticidal activity against the large milkweed bug (*Oncopeltus fasciatus*) and the Mediterranean fruit fly (*Ceratitis capitate*) (51). *P. crustosum* was also reported as the dominant species of the filamentous *Penicillium* spp. in extremely cold environments, in samples isolated from subglacial ice of arctic glaciers in Norway (52). Symbiotic relationships between fungi and their hosts are well established in three model organisms ranging from mutualism (*e.g.* fungus *Mortierella elongata* and algae *Nannochloropsis oceanica*) (53), commensalism (*e.g. Saccharomyces cerevisiae* or *Candida albicans* in the human digestive tract) (54) to parasitism (*e.g. Beauveria bassiana* in arthropods) (55). For instance, the pathogenic fungus *Beauveria bassiana* can utilise a cross-kingdom small-RNA effector to attenuate host immunity, thereby facilitating its infection in mosquito cells (56) or directly in insecticide-resistant *Ae. aegypti* mosquitoes (57). It would therefore be interesting to investigate further whether *Penicillium crustosum* and its mycotoxins affect mosquitoes’ life history traits, vector competence, or downstream viral infection in the mammalian host.

To date, there is a growing understanding of microbiota-arbovirus interactions and an expanding interest in using the microbiota for vector control and arbovirus outbreak containment. During a mosquito bite, saliva and virus are inserted simultaneously into the skin of the host. (58)Therefore, a thorough investigation into saliva microbiota and their potential contributions to critical stages in mosquito-borne virus infections is warranted. To investigate this, the presence of fungi in mosquito saliva needs to be eliminated. We obtained this by treating adults with the antifungal drug Fungin via the sugar meal. Depletion of the bacteria from the mosquito saliva was obtained using an oral treatment with an antibiotic cocktail consisting of gentamycin, penicillin, and streptomycin. Although this did not fully clear the bacteria from the entire mosquito body, we could obtain complete depletion of culturable bacteria in the mosquito saliva (**Fig 6**). Next, we aimed to investigate whether the saliva microbiota of *Ae. aegypti* interacts with viral particles in the mosquito saliva. Co-incubation of saliva from antibiotic or antifungal treated mosquitoes with SFV significantly reduced the infectious virus load in human cells (**Fig 7A**). Also pre-incubation of mosquito saliva from treated mosquitoes together with the virus before infecting cells lowered the amount of infectious virus (**Fig 7B**). In contrast, in both conditions, filtered saliva had no effect on viral infectivity compared to saliva from untreated control mosquitoes, suggesting that the physical presence of the saliva microbiota had no effect on infectivity. Moreover, the antiviral effect observed with saliva from AB treated mosquitoes is likely not due to presence of remaining bacterial secretions, since these secretions are also still present in the filtered saliva. We therefore hypothesize that the oral AB and/or AF treatment affected the composition of saliva components, beyond the microbial composition. Another option could be that the orally ingested AB or AF is carried over in the saliva, affecting virus infectivity.

In the cell pretreatment condition (**Fig 7C**), the incubation with NT saliva resulted in a downward trend in SFV titers, although this reduction was not significant. In contrast to the other incubation conditions, AB saliva did not result in a reduction in viral titers when cells were pretreated, whereas filtered saliva did reduce viral infectivity. This suggests that the viral inhibition could be due to an effect of bacterial secretions on the cells. The results of NT and AB saliva on pretreated cells are similar to prior research showing that mosquito saliva slightly diminished SFV replication in skin fibroblasts (59). The authors hypothesised that the presence of the saliva bacteria activated immune responses in the skin cells, leading to a higher viral resistance. However, in the pre-incubation with virus condition, we did not observe a similar reduction by mosquito saliva (from untreated *Ae. aegypti*) as seen in (59). This discrepancy could be due to the use of mosquitoes with different microbiota communities (*Ae. aegypti* Liverpool strain vs *Ae. aegypti* Pacific Paea strain) or different virus strains (SFV constructed from plasmid as per (60) vs SFV Vietnam strain). Moreover, our study used a lower MOI for infection (0.001 vs 0.01) and a longer pre-incubation period of virus with saliva (1h vs 20 min). Importantly, it has been equally shown that the virus-enhanced infection phenotype by mosquito saliva observed in mouse models could not be recapitulated *in vitro* or *ex vivo* (59). Therefore, it is imperative to conduct *in vivo* studies to elucidate the potential effects of saliva bacteria and fungi on virus infection and pathogenesis.

In conclusion, mosquito saliva fungi and bacteria were detected by culturing saliva from two distinct mosquito species: *Ae. aegypti* and *Cx. pipiens*. Furthermore, 16S metagenomics analysis showed that the bacterial community in saliva was more diverse than in midguts. The application of antibiotics or an antifungal drug in the oral treatment of adult mosquitoes depleted microbiota in the saliva, indicating that the origin of saliva microbiota lies within the mosquito body. Moreover, the depletion of fungi and bacteria from saliva may influence resistance to viral infection *in vitro*. However, the impacts of saliva microbiota warrant further investigation *in vivo*. Collectively, this work lays the foundation for further investigation into the effect of saliva fungi and bacteria on arbovirus replication and infection.

## Methods

### Ethics statement

Permits for mosquito field collections in Leuven were obtained from KU Leuven and the City Green Management (CGM) of Leuven. Mosquitoes of the *Ae. aegypti* lab strain were maintained by blood feeding on mice and were performed with the approval and under the guidelines of the Ethical Committee of the University of Leuven (license P053/2019).

### Mosquito strains

The *Ae. aegypti* Paea lab strain (provided by Pasteur Institute Paris via the EU-funded Infravec consortium) was reared and maintained under controlled conditions (28 ± 1°C, relative humidity of 80%, light: dark cycle of 16 h: 8 h). Field *Cx. pipiens* mosquitoes were collected in Leuven (N 50°52’41, E 4°41’21, Belgium) during the summer of 2021. Adult mosquitoes were trapped with BG-Sentinel traps (BioGents GmbH, Germany) with the help of BG-lure (BioGents GmbH, Germany). After sorting mosquitoes based on morphology, *Cx. pipiens* were maintained under conditions of 25 ± 1°C, relative humidity of 50%, and light: dark cycle of 14 h: 10 h. Adult mosquitoes were housed in Bugdorm-1 insect cages (30 x 30 x 30 cm, Bugdorm, Taiwan, China) and fed with 10% sterile sugar solution.

### Species identification of field mosquitoes by multiplex qPCR

Species identification was performed as described earlier (61). In brief, wings and legs of individual female mosquitoes were homogenized by bead disruption (2.8 mm Precellys, Bertin Instruments). Subsequently, the supernatant was hydrolysed at 95°C for 10 min. Multiplex qPCR was performed using a forward primer (5’-GCGGCCAAATATTGAGACTT-‘3) and reverse primer (5’-CGTCCTCAAACATCCAGACA-‘3) targeting the gene locus for acetylcholinesterase 2 to identify the *Cx. pipiens* complex(62). Additionally, subspecies-specific probes were used by targeting the CQ11 microsatellite locus to distinguish between *Cx. pipiens pipiens* (5’-GCTTCGGTGAAGGTTTGTGT-’3)(63) and *Cx. pipiens molestus* (5’-TGAACCCTCCAGTAAGGTATCAACTAC-‘3)(64). Reactions were prepared in a final volume of 25 µL containing 3 µL of template DNA, 6.25 µL of 2x TaqMan™ PCR Master mix, 0.38 µL of each primer at 10 µM, 1 µL of each probe at 5 µM, 0.06 µL Reverse Transcriptase and 12.94 µL of nuclease-free water. The thermal profile consisted of an initial step of 15 min at 95°C, followed by 40 amplification cycles of 1 min at 94°C, 1 min at 48°C and 1 min at 72°C.

### Forced salivation and dissection of midguts and salivary glands

Mosquitoes were starved 24h prior to salivation. Female *Ae. aegypti* or *Cx. pipiens* mosquitoes were anaesthetized by placing them under cold conditions (4-8°C) for 5 min, or by CO_2_ released by dry ice for 2 min, respectively. While sedated, legs and wings were removed to immobilize the mosquito during salivation. Saliva for culturing was harvested by gently placing the proboscis of a living female into a 20 µL pipette tip (uTIP, Biotix) containing 20 µL of a 1:1 solution of 10% sterile sugar solution and 1X PBS. Saliva for the *in vitro* infection assays was harvested by gently placing the proboscis of a living female into the pipette tip containing 1 µL of silicon oil as described previously (59). Blank tips containing only the salivation solution were placed aside on the bench as the environment control during each salivation. After 1h of forced salivation, saliva was harvested and stored at -80°C either individually or pooled for further studies.

The surface of the mosquito body was disinfected individually by 75% ethanol and rinsed twice with 1X PBS buffer before dissection of the salivary glands and midgut. Salivary glands were harvested before the midgut to avoid potential contamination as the midgut harbours the highest bacterial load. The salivary glands and midguts were pooled (8 to 12 females) in homogenization tubes (2.8 mm Precellys, Bertin Instruments) containing 150-200 µL of 1X PBS and then stored at -80°C for further analysis.

### Artificial blood feeding, antibiotic and antifungal treatment

To study the effects of a blood meal on saliva microbiota, 5 to 7-day-old *Ae. aegypti* female mosquitoes were orally fed for 30 min using an artificial membrane feeding system (Hemotek, UK). The bloodmeal contained fresh rabbit erythrocytes washed with PBS and plasma was replaced by the same volume of sterile FBS. Fully engorged females were cold-anesthetized and separated to be maintained up to two weeks until forced salivation, under the same controlled conditions mentioned above. In parallel, the blood meal was cultured on Luria-Bertani (LB) plates to determine the absence of bacteria in the blood meal.

Adult mosquitoes were treated with an antibiotic cocktail with doses ranging from 10 units/mL penicillin and 10 µg/mL streptomycin/gentamycin (Penicillin-Streptomycin 100X, Gentamycin, Gibco™) up to 200 units/mL penicillin and 200 µg/mL streptomycin/gentamycin into the 10% sugar solution for 6 days. For the antifungal treatment, 25 µg/mL of Fungin™ (InvivoGen) was added into the 10% sugar solution for 10 days. The sugar solution containing antibiotic and/or antifungal compounds was refreshed every day throughout the treatment.

### Bacteria and fungi culturing on agar plates

Mosquito-derived samples (saliva, salivary glands, and midguts) were cultured on LB broth, Brain Heart Infusion (BHI) broth or blood agar plates at 28°C to compare microbial growth on different media. Saliva from 8-12 mosquitoes was pooled. Approximately 50 µL of the saliva pool was inoculated per agar plate. For the salivary gland samples, 50 µL of supernatant post-homogenization was used. For the midgut, the supernatant was diluted 10 to 100 times in sterile water post-homogenization before culturing 50 µL of supernatant on the agar plates. As a negative control, the salivation solution was cultured in parallel. Plates were checked after 24, 48 and 120 h of incubation for bacteria, and up to one month for fungi. Colony-forming units (CFU) were calculated to determine the bacterial and fungal loads.

### Bacterial identification by 16S Sanger sequencing

Individual colonies were selected from the agar plates and resuspended in 30 µL of sterile water. The bacterial suspension was heat-inactivated at 95°C for 15 min. When necessary, the DNA concentration was diluted to 100 ng/µL. PCR amplification of 467 bp in the 16S ribosomal RNA gene was performed with the 16S forward universal primer (5’ TCC TAC GGG AGG CAG CAG T ’3) and 16S reverse universal primer (5’ GGA CTA CCA GGG TAT CTA ATC CTG TT ’3) as described previously by Romoli et al(65). The PCR product was confirmed by gel electrophoresis (1% agarose gel). PCR samples were purified by the Wizard SV Gel and PCR Clean system following manufacturers’ instructions. After purification, samples were sent to Macrogen Europe for Sanger sequencing. The DNA sequences obtained were trimmed using TrimAl (v. 1.3) and searched for homology to those available at the GenBank-EMBL database using the Basic Local Alignment Search Tool (BLAST) program (NCBI).

### Fungal identification by ITS3-ITS4 and CRU-specific Sanger sequencing

A portion of the fungal colony was collected into a homogenization tube with 500 µL of PBS. Homogenization was performed at 2 cycles of 6800 rpm for 10 sec with a pause of 20 sec between each cycle. After homogenization, 200 µL of the supernatant was used for DNA extraction, which was performed using the QIAamp DNA Kit 250 (51306, QIAGEN) following manufacturers’ instructions. The universal forward primer targeting the ITS3 region (5’ GCA TCG ATG AAG AAC GCA GC ’3) and universal reverse primer targeting the ITS4 region (5’ TCC TCC GCT TAT TGA TAT GC ’3) were used for ITS PCR, amplifying a 350 bp PCR product as described previously(27). For species-specific detection of *Penicillium crustosum*, the CRU-F forward primer (5’ TCC CAC CCG TGT TTA TTT TA ‘3) and CRU-R reverse primer (5’ TCC CTT TCA ACA ATT TCA CG ‘3) were used, amplifying a 892 bp PCR product(66). The length of the PCR products was confirmed by gel electrophoresis (1% agarose gel). PCR samples were purified by the Wizard SV Gel and PCR Clean system (Promega) following manufacturers’ instructions. The protocol for Sanger sequencing and analysis is identical as described above for the bacterial identification.

### DNA extraction from midgut, salivary glands, and saliva samples

DNA extractions were strictly performed under a laminar flow cabinet. Blank Eppendorf tubes were added per experiment as a negative environmental control. DNA extractions were performed using the QIAamp DNA Kit 250 (51306, QIAGEN) following an adapted version of the manufacturers’ protocol including an additional step of adding 0.5 µL of Ready-Lyse™ Lysozyme Solution (Lucigen) to enable the collection of DNA from both Gram+ and Gram-species. Next, 100 µL ATL buffer and 20 µL proteinase K were added to each sample, followed by an incubation step at 56°C for 5 min. After pulse-vortexing for 15 sec, samples were incubated at 70°C for 10 min and 230 µL of ethanol was added. The following steps followed the original protocol according to manufacturers’ instructions. Samples were eluted in 50 µL AE buffer and were stored at -20°C until further analysis.

### Quantification of bacterial and fungal loads by qPCR

The bacterial and fungal loads in extracted DNA samples were quantified by qPCR using the iTaq Universal SYBR Green One-Step kit (Bio-Rad). Each reaction was prepared in a final volume of 20 µL containing 5 µL of template DNA, 10 µL of 2x one-step SYBR Green reaction mix, 0.4 µL of each primer at 10 µM and 4.2 µL of nuclease-free water. The same primers were used as for Sanger sequencing (described above). The thermal profile consisted of an initial step of 10 min at 95°C, followed by 40 amplification cycles of 15 sec at 95°C and 1 min at 50°C for bacteria, while a 45°C annealing temperature was used for fungal quantification. The genome copies/mosquito for each sample were quantified based on the use of a gBlocks™ Gene Fragment (IDT) as a standard dilution for both the 16S and ITS qPCR. Limit of detection (LOD) was determined as the lowest bacterial/fungal load able to be detected by the qPCR assay in 95% of experiments.

### Phylogenetic analysis

Sanger sequencing was performed by Macrogen Europe. The consensus sequence of fungal or bacterial colonies together with reference sequences from GenBank were aligned with MEGA-X. The resulting alignment was trimmed by using TrimAl (v.1.3). A maximum likelihood (ML) tree was constructed using MEGA-X with the substitution model being Kamura 2-parameter model (K2) and Gamma distributed (G). 500 bootstrap replicates were used. Finally, trees were visualized using FigTree v1.4.4.

### 16S metagenomic sequencing analysis

DNA extractions were performed using the QIAamp DNA Kit 250, including an additional step of adding Ready-Lyse™ Lysozyme Solution as described above. Subsequently, the DNA was sent to Macrogen company to perform the following pipeline: Library prep (16S rRNA V3-V4 Illumina) and Sequencing (MiSeq - 300 paired-end; 200K reads/100K paired-end reads). High-quality sequences were obtained from Macrogen. Raw data were analyzed via the DADA2 pipeline. In brief, paired-end sequences were checked, de-replicated, filtered and merged, and chimera sequences were removed (consensus) using the DADA2 quality control method. The putative error-free sequences were referred to as sequence variants (SVs). Sequenced samples were provided as amplicon sequence variant (ASV) abundance tables for further analysis. SVs were assigned taxonomy using the assignTaxonomy function within DADA2 with the naive Bayesian classifier method, trained on the Silva version138 99% database, into phylum, class, order, family, and genus levels to evaluate the corresponding taxonomic abundance.

### Virus strains and cells

The SFV Vietnam strain (Genbank EU350586.1) belongs to the collection of the Rega Institute for Medical Research (Belgium). Virus stocks were propagated in *Ae. albopictus-* derived C6/36 cells (ATCC CRL-1660) at 28°C and stored at -80°C. Viral titers were determined by endpoint titration and plaque assay on Vero cells. African green monkey kidney epithelial cells (Vero cells, ATCC CCL-81) were maintained in minimal essential medium (MEM, Gibco) supplemented with 10% fetal bovine serum (FBS, Gibco), 1% L-glutamine (L-glu, Gibco), 1% sodium bicarbonate (NaHCO_2_, Gibco), and 1% non-essential amino acids (NEAA, Gibco). Human skin fibroblasts (ATCC CRL-2522) were grown in MEM supplemented with 10% FBS, 1% L-glu, 1% NaHCO_2_, 1% NEAA and 1% sodium pyruvate (Gibco). Cell cultures were maintained at 37°C in an atmosphere of 5% CO_2_ and 95%-99% relative humidity. Assays were performed using a similar medium but supplemented with 2% FBS (2% assay medium).

### *In vitro* assays of co-incubation of mosquito saliva with Semliki Forest virus

Human skin fibroblast cells were pre-seeded (12 000 cells/well, in 2% assay medium) in 96 well-plates 24h before the infection. Saliva from non-treated *Ae. aegypti*, AF saliva from antifungal-treated females, AB saliva from antibiotic-treated females, and filtered saliva from non-treated *Ae. aegypti* using a 0.2 µm filter were assessed. Three scenarios were evaluated: (1) cells were directly inoculated with saliva and virus (MOI:0.001) for 1h; (2) virus was pre-incubated with saliva for 1h at 28°C before being added to the cells for 1h (MOI:0.001); (3) cells were preincubated with saliva for 1h followed by virus infection (MOI: 0.001) for 1h. After virus infection for 1h, the virus inoculum was removed, cells were washed 3X with PBS and subsequently, a fresh 2% assay medium was added. The plates were maintained at 37°C with 5% of CO_2_ until 24h post-infection, when supernatant was harvested and stored at -80°C for further analysis. The levels of infectious virus in the supernatant were determined by end-point titrations on Vero cells. Cells were pre-seeded in 96 well-plates (25 000 cells/well) in 2% assay medium and incubated overnight (37°C, 5% CO_2_). The next day, 10-fold serial dilutions of the samples were prepared in triplicate in the plates and incubated for 3 days (37°C, 5% CO_2_). At day 3 pi, cells were scored microscopically for virus-induced cytopathogenic effect (CPE). The tissue culture infectious dose_50_/ml (TCID_50_/ml) was calculated using the Reed and Muench method(67). Limit of quantification (LOQ) was determined as the lowest TCID_50_ value that could be quantified by the end-point titration assay.

### Statistical analysis

GraphPad Prism 9.4.1 and R (version 4.1.0) were used to make graphs and perform statistical analyses. In DADA2, alpha diversity in the microbial communities identified from different organs was visualised by Phyloseq and scored using the “observed” and “Shannon” diversity indexes and Kruskal-Wallis tests were used. Beta diversity was assessed using Bray-Curtis distances. The bacterial and fungal loads were statistically analysed using the Mann-Whitney test (two groups) or the Kruskal-Wallis test (≥ three groups). Statistical details are described in the figure legends. The statistical significance threshold was assessed at p < 0.05.

## Acknowledgements

This project was funded by KU Leuven (C22/18/007, C14/20/108 and starting grant STG/19/008). This work was also supported by the project “Research Infrastructures for the control of vector-borne diseases” (Infravec2), which has received funding from the European Union’s Horizon 2020 research and innovation program under grant agreement number 731060. We would like to thank the City Green Management of Leuven, Belgium, for providing the permits for the mosquito field collections during the summer of 2021. We appreciated the help of Dr. Daniella A. Lefteri (University of Leeds) for sharing the protocol for forced salivation with oil. We also appreciated the help of Prof. Mutien Garigliany and Felipe Rivas from Université de Liège, for testing the saliva of *Culex* mosquitoes in vitro.

## Competing interests

The authors declare no competing interests.

## Author contributions

Conceptualization and Methodology: LW, LR, KT, LD; Investigation: LW, LR, NA; Data analysis: LW, LR, NA, KT, LD; Writing – Original Draft Preparation: LW, KT, LR, AS, SW, JB, MG, LD.

## Data availability

The *Penicillium crustosum* sequence identified in this study has been deposited in GenBank under the accession number OL436123. Additionally, bacterial sequences retrieved in this study have been deposited in GenBank under the accession numbers: *Asaia* sp. ON527783; *Chryseobacterium* sp. ON527789; *Enterobacter* sp. ON527791; *Microbacterium* sp. ON527869; *Klebsiella* sp. ON527877; *Serratia nematodiphila* ON527870; *Serratia marcescens* ON527875. The other sequences used for the analysis were available from GenBank. NGS raw data for *Ae. aegypti* mosquito saliva microbiota 16S rRNA V3–V4 metagenomic sequencing data was submitted in Mendeley Data with DOI: 10.17632/mvd793vm6v.1 (active when the paper gets published).

## Notes

### Competing Interest Statement

The authors have declared no competing interest.

### Summary of Updates

New analyses were performed, and extra results were added.

## References

1. Tandina F, Doumbo O, Yaro AS, Traoré SF, Parola P, Robert V. Mosquitoes (Diptera: Culicidae) and mosquito-borne diseases in Mali, West Africa.

2. Gould E, Pettersson J, Higgs S, Charrel R, de Lamballerie X. Emerging arboviruses: Why today? One Health. 2017 Dec 1;4:1.

3. Shragai T, Tesla B, Murdock C, Harrington LC. Zika and chikungunya: mosquito-borne viruses in a changing world. Ann N Y Acad Sci. 2017;1399(1):61–77.

4. Pang X, Zhang R, Cheng G. Progress towards understanding the pathogenesis of dengue hemorrhagic fever. Virol Sin. 2017 Feb 1;32(1):16–22.

5. Colpitts TM, Conway MJ, Montgomery RR, Fikrig E. West Nile Virus: biology, transmission, and human infection. Clin Microbiol Rev. 2012 Oct;25(4):635–48.

6. Achee NL, Grieco JP, Vatandoost H, Seixas G, Pinto J, Ching-Ng L, et al. Alternative strategies for mosquito-borne arbovirus control. PLoS Negl Trop Dis. 2019;13(1).

7. World Health Organization. Global vector control response 2017–2030. World Health Organization. 2017. 52 p.

8. Corbel V, Achee NL, Chandre F, Coulibaly MB, Dusfour I, Fonseca DM, et al. Tracking Insecticide Resistance in Mosquito Vectors of Arboviruses: The Worldwide Insecticide resistance Network (WIN). PLoS Negl Trop Dis. 2016 Dec 1;10(12).

9. Montenegro D, Cortés-Cortés G, Balbuena-Alonso MG, Warner C, Camps M. Wolbachia-based emerging strategies for control of vector-transmitted disease. Vol. 260, Acta Tropica. Elsevier B.V.; 2024.

10. Wilke ABB, Marrelli MT. Paratransgenesis: A promising new strategy for mosquito vector control. Vol. 8, Parasites and Vectors. BioMed Central Ltd.; 2015.

11. Franz AWE, Kantor AM, Passarelli AL, Clem RJ. Tissue Barriers to Arbovirus Infection in Mosquitoes. Viruses. 2015 Jul 8;7(7):3741.

12. Agarwal A, Parida M, Dash PK. Impact of transmission cycles and vector competence on global expansion and emergence of arboviruses. Rev Med Virol. 2017 Sep 1;27(5):e1941.

13. Mancini M V., Damiani C, Accoti A, Tallarita M, Nunzi E, Cappelli A, et al. Estimating bacteria diversity in different organs of nine species of mosquito by next generation sequencing. BMC Microbiol. 2018 Oct 4;18(1):1–10.

14. Kang X, Wang Y, Li S, Sun X, Lu X, Rajaofera MJN, et al. Comparative Analysis of the Gut Microbiota of Adult Mosquitoes From Eight Locations in Hainan, China. Front Cell Infect Microbiol. 2020 Dec 15;10:791.

15. Bozic J, Capone A, Pediconi D, Mensah P, Cappelli A, Valzano M, et al. Mosquitoes can harbour yeasts of clinical significance and contribute to their environmental dissemination. Environ Microbiol Rep. 2017 Oct 1;9(5):642–8.

16. Gusmão DS, Santos A V., Marini DC, Bacci M, Berbert-Molina MA, Lemos FJA. Culture-dependent and culture-independent characterization of microorganisms associated with Aedes aegypti (Diptera: Culicidae) (L.) and dynamics of bacterial colonization in the midgut. Acta Trop. 2010 Sep;115(3):275–81.

17. Dennison NJ, Jupatanakul N, Dimopoulos G. The mosquito microbiota influences vector competence for human pathogens. Curr Opin Insect Sci [Internet]. 2014;3:6–13. Available from: http://linkinghub.elsevier.com/retrieve/pii/S221457451400039X

18. Apte-Deshpande AD, Paingankar MS, Gokhale MD, Deobagkar DN. Serratia odorifera mediated enhancement in susceptibility of Aedes aegypti for chikungunya virus. Indian J Med Res. 2014;139(5):762–8.

19. Apte-Deshpande A, Paingankar M, Gokhale MD, Deobagkar DN. Serratia odorifera a midgut inhabitant of Aedes aegypti mosquito enhances its susceptibility to dengue-2 virus. PLoS One. 2012 Jul 27;7(7).

20. Wu P, Sun P, Nie K, Zhu Y, Shi M, Xiao C, et al. A Gut Commensal Bacterium Promotes Mosquito Permissiveness to Arboviruses. Cell Host Microbe. 2019 Jan 9;25(1):101–112.e5.

21. Angleró-Rodríguez YI, Talyuli OAC, Blumberg BJ, Kang S, Demby C, Shields A, et al. An Aedes aegypti-associated fungus increases susceptibility to dengue virus by modulating gut trypsin activity. Elife. 2017 Dec 5;6.

22. Conway MJ, Colpitts TM, Fikrig E. Role of the Vector in Arbovirus Transmission.

23. Manning JE, Morens DM, Kamhawi S, Valenzuela JG, Memoli M. Mosquito Saliva: The Hope for a Universal Arbovirus Vaccine? J Infect Dis. 2018 Jun 5;218(1):7.

24. Demarta-Gatsi C, Mécheri S. Vector saliva controlled inflammatory response of the host may represent the Achilles heel during pathogen transmission. J Venom Anim Toxins Incl Trop Dis. 2021 May 1;27:20200155.

25. Accoti A, Damiani C, Nunzi E, Cappelli A, Iacomelli G, Monacchia G, et al. Anopheline mosquito saliva contains bacteria that are transferred to a mammalian host through blood feeding. Front Microbiol. 2023;14.

26. Onyango MG, Payne AF, Stout J, Dieme C, Kuo L, Kramer LD, et al. Aedes albopictus saliva contains a richer microbial community than the midgut. Parasit Vectors. 2024 Dec 1;17(1).

27. Maza-Márquez P, Vílchez-Vargas R, González-Martínez A, González-López J, Rodelas B. Assessing the abundance of fungal populations in a full-scale membrane bioreactor (MBR) treating urban wastewater by using quantitative PCR (qPCR). J Environ Manage. 2018 Oct;223:1–8.

28. Demjanová S, Jevinová P, Pipová M, Regecová I. Identification of Penicillium verrucosum, Penicillium commune, and Penicillium crustosum Isolated from Chicken Eggs. Processes 2021, Vol 9, Page 53. 2020 Dec 29;9(1):53.

29. Badran RAM, Aly MZY. Studies on the mycotic inhabitants of Culex pipiens collected from fresh water ponds in Egypt. Mycopathologia. 1995 Nov;132(2):105–10.

30. Moosa-Kazemi SH, Asgarian TS, Sedaghat MM, Javar S. Pathogenic fungi infection attributes of malarial vectors Anopheles maculipennis and Anopheles superpictus in central Iran. Malar J. 2021 Dec 1;20(1):393.

31. Gusmão DS, Santos A V., Marini DC, Bacci M, Berbert-Molina MA, Lemos FJA. Culture-dependent and culture-independent characterization of microorganisms associated with Aedes aegypti (Diptera: Culicidae) (L.) and dynamics of bacterial colonization in the midgut. Acta Trop. 2010 Sep;115(3):275–81.

32. Chouaia B, Rossi P, Montagna M, Ricci I, Crotti E, Damiani C, et al. Molecular Evidence for Multiple Infections as Revealed by Typing of Asaia Bacterial Symbionts of Four Mosquito Species. Appl Environ Microbiol. 2010;76(22):7444–50.

33. Favia G, Ricci I, Marzorati M, Negri I, Alma A, Sacchi L, et al. Bacteria of the genus Asaia: a potential paratransgenic weapon against malaria. Adv Exp Med Biol. 2008;627:49–59.

34. Kang X, Wang Y, Li S, Sun X, Lu X, Rajaofera MJN, et al. Comparative Analysis of the Gut Microbiota of Adult Mosquitoes From Eight Locations in Hainan, China. Front Cell Infect Microbiol. 2020 Dec 15;10:791.

35. Coon KL, Brown MR, Strand MR. Gut bacteria differentially affect egg production in the anautogenous mosquito Aedes aegypti and facultatively autogenous mosquito Aedes atropalpus (Diptera: Culicidae). Parasit Vectors. 2016 Jun 30;9(1).

36. Coon KL, Vogel KJ, Brown MR, Strand MR. Mosquitoes rely on their gut microbiota for development. Mol Ecol. 2014;23(11):2727–39.

37. Hyde J, Gorham C, Brackney DE, Steven B. Antibiotic resistant bacteria and commensal fungi are common and conserved in the mosquito microbiome. PLoS One. 2019 Aug 1;14(8):e0218907.

38. Accoti A, Damiani C, Nunzi E, Cappelli A, Iacomelli G, Monacchia G, et al. Anopheline mosquito saliva contains bacteria that are transferred to a mammalian host through blood feeding. Front Microbiol. 2023 Jul 18;14:1157613.

39. Onyango MG, Payne AF, Stout J, Dieme C, Kuo L, Kramer LD, et al. Aedes albopictus saliva contains a richer microbial community than the midgut. Parasit Vectors. 2024 Dec 1;17(1):267.

40. Gómez MMDLNVLYPYCBOCMPLHMMRJD. Comparative analysis of bacterial microbiota in Aedes aegypti (Diptera: Culicidae): insights from field and laboratory populations in Colombia. J Med Entomol. 2025;62(2):358–70.

41. Malassigné S, Moro CV, Luis P. Mosquito Mycobiota: An Overview of Non-Entomopathogenic Fungal Interactions. Pathogens. 2020 Jul 1;9(7):564.

42. Malassigné S, Moro CV, Luis P. Mosquito mycobiota: An overview of non-entomopathogenic fungal interactions. Vol. 9, Pathogens. MDPI AG; 2020. p. 1–14.

43. Da Costa GL, De Oliveira PC. Penicillium species in mosquitoes from two Brazilian regions. J Basic Microbiol. 1998;38(5–6):343–7.

44. Moosa-Kazemi SH, Asgarian TS, Sedaghat MM, Javar S. Pathogenic fungi infection attributes of malarial vectors Anopheles maculipennis and Anopheles superpictus in central Iran. Malar J. 2021 Dec 1;20(1).

45. Da E, Pereira S, De M Sarquis MI, Ferreira-Keppler RL, Hamada N, Alencar YB. Filamentous Fungi Associated with Mosquito Larvae (Diptera: Culicidae) in Municipalities of the Brazilian Amazon. ECOLOGY, BEHAVIOR AND BIONOMICS.

46. Badran RAAM. Studies on the mycotic inhabitants ofCulex pipiens collected from fresh water ponds in Egypt. Human And Animal Mycology. 1995;132:105–10.

47. Angleró-Rodríguez YI, Blumberg BJ, Dong Y, Sandiford SL, Pike A, Clayton AM, et al. A natural Anopheles-associated Penicillium chrysogenum enhances mosquito susceptibility to Plasmodium infection. Sci Rep. 2016 Sep 28;6.

48. De Souza GG, Pfenning LH, De Moura F, Salgado M, Takahashi JA. Isolation, identification and antimicrobial activity of propolis-associated fungi. Nat Prod Res. 2013;27(18):1705– 7.

49. Rundberget T, Skaar I, Flåøyen A. The presence of Penicillium and Penicillium mycotoxins in food wastes. Int J Food Microbiol. 2004 Jan 15;90(2):181–8.

50. Moldes-Anaya A, Rundberget T, Uhlig S, Rise F, Wilkins AL. Isolation and structure elucidation of secopenitrem D, an indole alkaloid from Penicillium crustosum Thom. Toxicon. 2011 Feb 1;57(2):259–65.

51. Carmen González M, Lull C, Moya P, Ayala I, Primo J, Yúfera Primo E. Insecticidal Activity of Penitrems, Including Penitrem G, a New Member of the Family Isolated from Penicillium crustosum. J Agric Food Chem. 2003 Apr 9;51(8):2156–60.

52. Sonjak S, Frisvad JC, Gunde-Cimerman N. Genetic variation among Penicillium crustosum isolates from Arctic and other ecological niches. Microb Ecol. 2007 Aug;54(2):298–305.

53. Bonfante P. Algae and fungi move from the past to the future. Elife. 2019 Jul 1;8:e49448.

54. Tong Y, Tang J. Candida albicans infection and intestinal immunity. Microbiol Res. 2017 May 1;198:27–35.

55. Mascarin GM, Jaronski ST. The production and uses of Beauveria bassiana as a microbial insecticide. World Journal of Microbiology and Biotechnology 2016 32:11. 2016 Sep 15;32(11):1–26.

56. Cui C, Wang Y, Liu J, Zhao J, Sun P, Wang S. A fungal pathogen deploys a small silencing RNA that attenuates mosquito immunity and facilitates infection. Nat Commun. 2019 Dec 1;10(1):4298.

57. Cui C, Wang Y, Li Y, Sun P, Jiang J, Zhou H, et al. Expression of mosquito miRNAs in entomopathogenic fungus induces pathogen-mediated host RNA interference and increases fungal efficacy. Cell Rep. 2022 Oct 25;41(4):111527.

58. Scolari F, Casiraghi M, Bonizzoni M. Aedes spp. and Their Microbiota: A Review. Front Microbiol. 2019 Sep 4;10.

59. Lefteri DA, Bryden SR, Pingen M, Terry S, Mccafferty A, Beswick EF, et al. Mosquito saliva enhances virus infection through sialokinin-dependent vascular leakage. 2022; Available from: http://www.pnas.org/lookup/suppl/doi:10.1073/pnas.

60. Ülper L, Sarand I, Rausalu K, Merits A. Construction, properties, and potential application of infectious plasmids containing Semliki Forest virus full-length cDNA with an inserted intron. J Virol Methods. 2008 Mar;148(1–2):265–70.

61. Wang L, Soto A, Remue L, Rosales Rosas AL, De Coninck L, Verwimp S, et al. First Report of Mutations Associated With Pyrethroid (L1014F) and Organophosphate (G119S) Resistance in Belgian Culex (Diptera: Culicidae) Mosquitoes. J Med Entomol. 2022 Nov 16;59(6):2072–9.

62. 62. Rudolf M, Czajka C, Bö Rstler J, Melaun C, Jö St H, Von Thien H, et al. First Nationwide Surveillance of Culex pipiens Complex and Culex torrentium Mosquitoes Demonstrated the Presence of Culex pipiens Biotype pipiens/molestus Hybrids in Germany. 2013;

63. Rudolf M, Czajka C, Bö Rstler J, Melaun C, Jö St H, Von Thien H, et al. First Nationwide Surveillance of Culex pipiens Complex and Culex torrentium Mosquitoes Demonstrated the Presence of Culex pipiens Biotype pipiens/molestus Hybrids in Germany. 2013;

64. Vogels CBF, Van De Peppel LJJ, Van Vliet AJH, Westenberg M, Ibañez-Justicia A, Stroo A, et al. Winter activity and aboveground hybridization between the two biotypes of the west nile virus vector culex pipiens. Vector-Borne and Zoonotic Diseases. 2015 Oct;15(10):619–26.

65. Romoli O, Schönbeck JC, Hapfelmeier S, Gendrin M. Production of germ-free mosquitoes via transient colonisation allows stage-specific investigation of host–microbiota interactions. Nat Commun. 2021 Feb 11;12(1):1–16.

66. Demjanová S, Jevinová P, Pipová M, Regecová I. Identification of Penicillium verrucosum, Penicillium commune, and Penicillium crustosum Isolated from Chicken Eggs. Processes 2021, Vol 9, Page 53. 2020 Dec 29;9(1):53.

67. Reed LJ, Muench H. A simple method of estimating fifty per cent endpoints. Am J Epidemiol. 1938 May 1;27(3):493–7.

