## Supplementrary file for "Identification of a Culturable Fungal Species and Endosymbiotic Bacteria in Saliva of *Aedes aegypti* and *Culex pipiens* and Their Impact on Arbovirus Infection *in Vitro*"

### Supplementary Information

### Supplementary Figures

**Fig S1. Gel electrophoresis after PCR amplification.** DNA fragments were amplified with ITS3-4 (ITS) universal primers (350 bp) and a *Penicillium crustosum* specific primer pair (892 bp) on 3 fungal colonies tested.

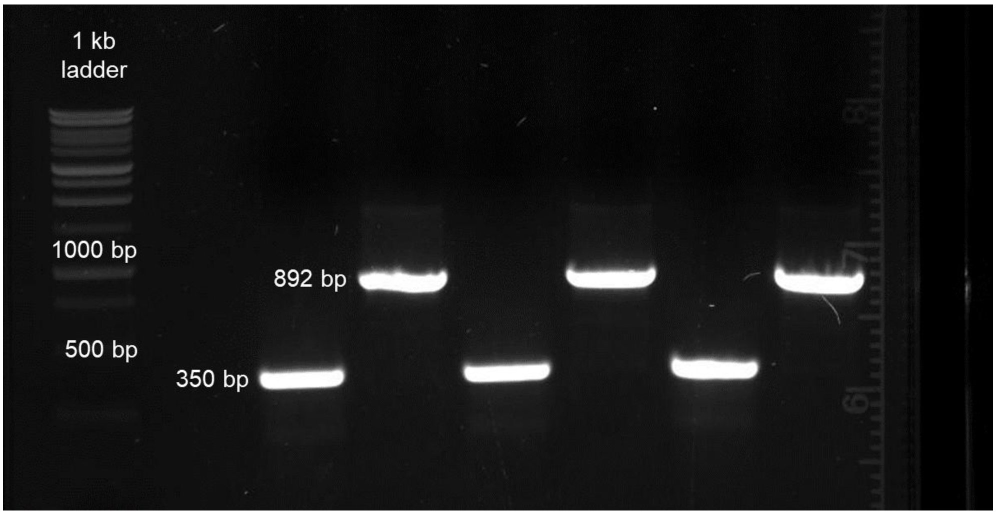

**Fig S2. CFU of salivary bacteria by culturing on different agar plates.** (A-B) Pooled saliva of *Aedes aegypti* cultured on LB, BHI and blood agar plates and incubated at 28°C under aerobic conditions. After 48 hours, colonies were counted, and CFU/mosquito was calculated for each agar plate for saliva from *Aedes aegypti*. Significant differences were demonstrated by the Kruskal-Wallis test (ns, p>0.05).

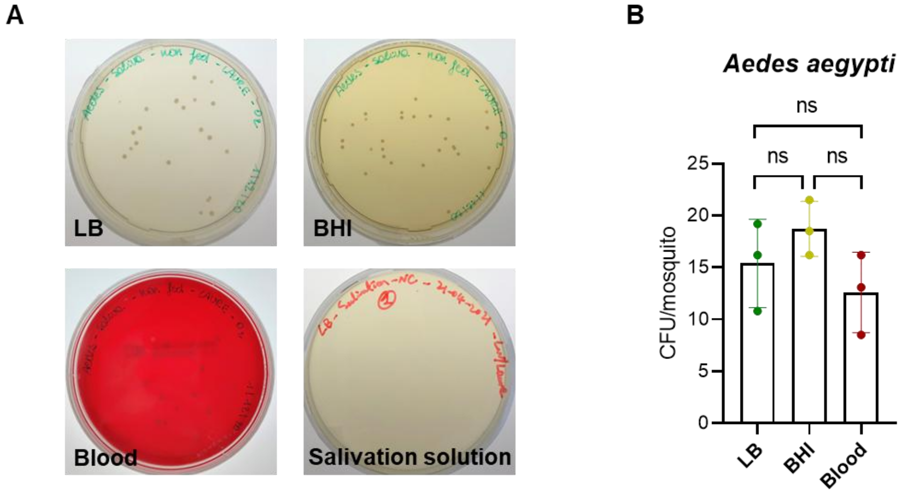

**Fig S3. Bacterial loads of individual midgut and salivary glands of *Culex pipiens* and *Aedes aegypti* mosquitoes.** The CFU read-out was performed by visual counting after culturing the homogenized tissues for 48h at 28°C on LB agar plates. Negative control (NC) consisted of the blank solution used as environment control for forced salivation. For the salivary gland samples, 50 µL of supernatant post-homogenization was used. For the midgut, the supernatant was diluted 10 to 100 times in sterile water post-homogenization before culturing 50 µL of the supernatant on the agar plates. SG = salivary glands. The lines represent the median values.

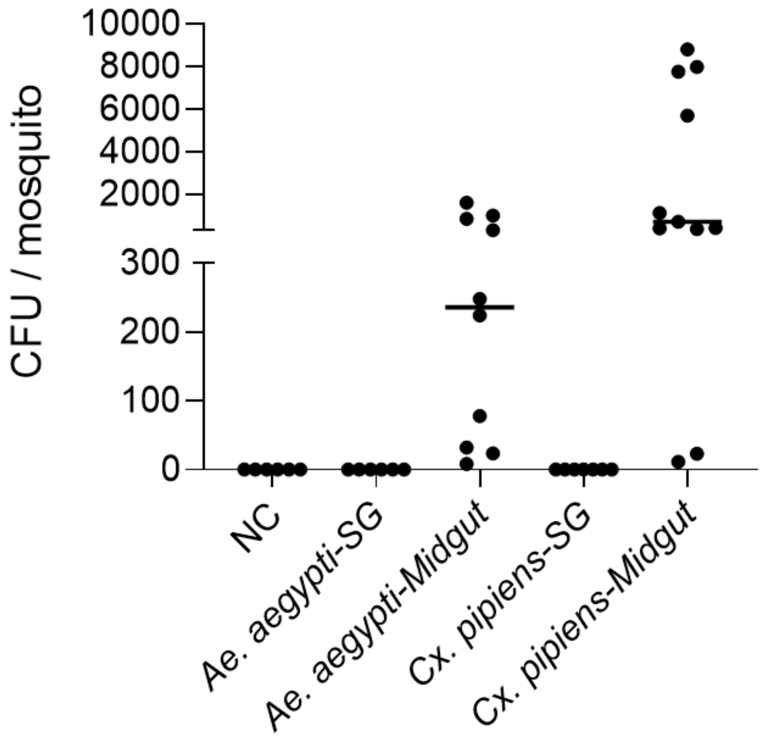

**Fig S4. Bacterial loads in different life stages of *Aedes aegypti*.** The CFU read-out was performed by visual counting after culturing the homogenized mosquito bodies for 48h at 28°C on LB agar plates. The lines represent the median values. D0: newly emerged adults, Negative control (NC) consisted of the blank solution used as environment control for forced salivation. Statistical analysis was done using the Kruskal-Wallis test (*, p<0.05; **, p<0.01).

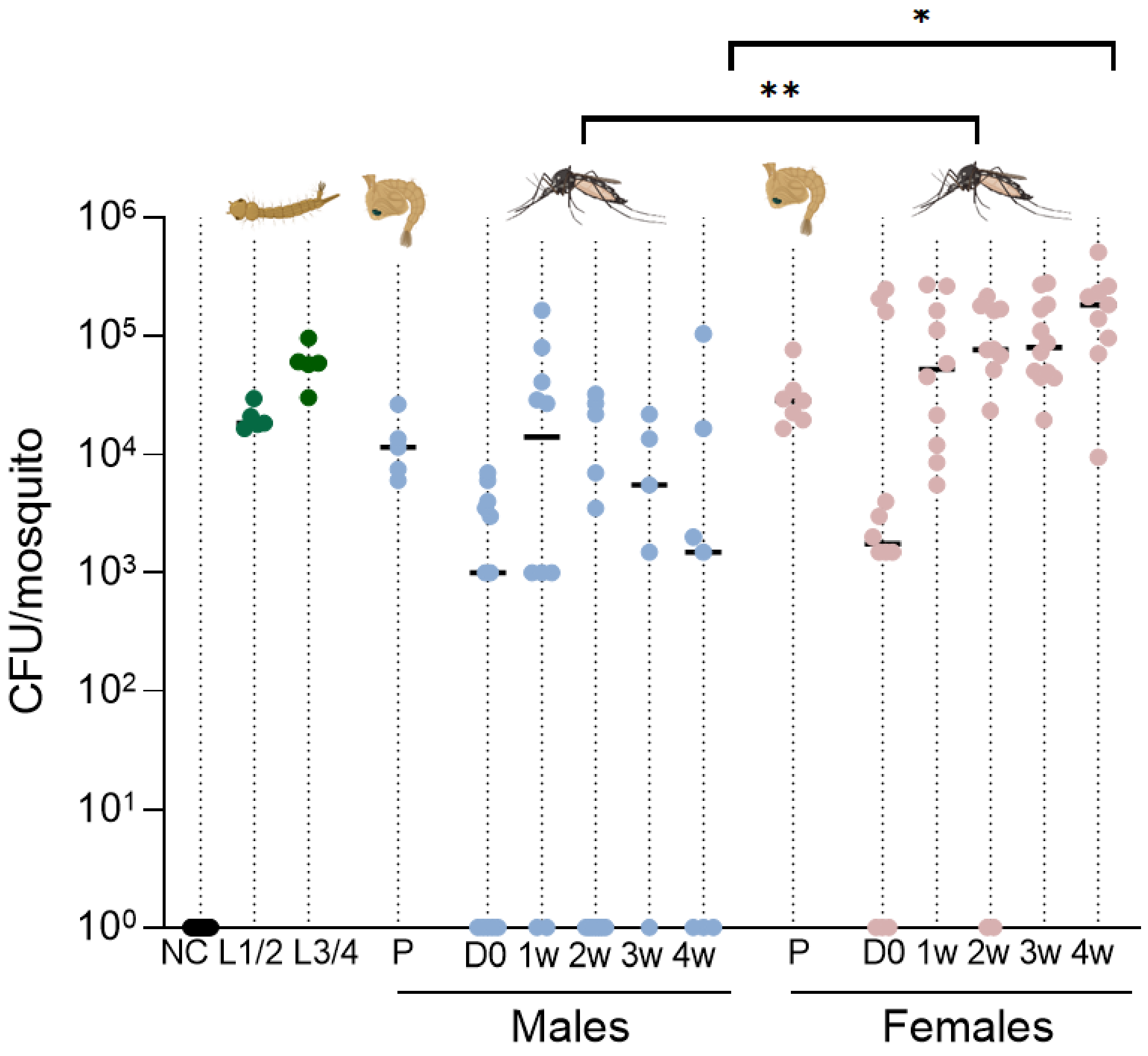

**Fig S5. Bacteria composition in mosquitoes variation according to life stage and sex.** Based on the 16S Sanger sequencing, number of colonies (%) per genus was calculated among pupae males; newly emerged males (D0); 2-week-old males; pupae females; newly emerged females (D0); and 2-week-old females.

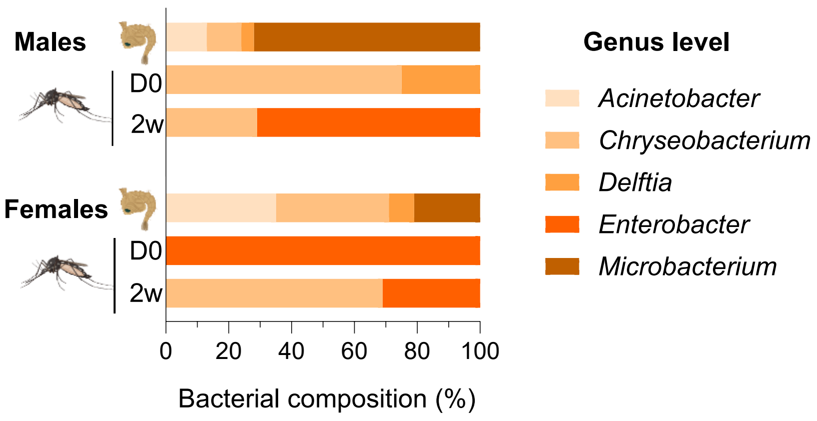

**Fig S6.** Survival rate of mosquitoes recorded for each treatment group. Female *Aedes aegypti* adults (5-7 days old) were divided between three treatment groups: untreated control group (NT: only 10% sucrose), antifungal-treated (AF: 25 µg/mL of Fungin in 10% sucrose for 10 days) and antibiotic-treated (AB: serial dilution up to 200 units/mL of penicillin and 200 µg/mL streptomycin + gentamycin in 10% sucrose for 6 days).

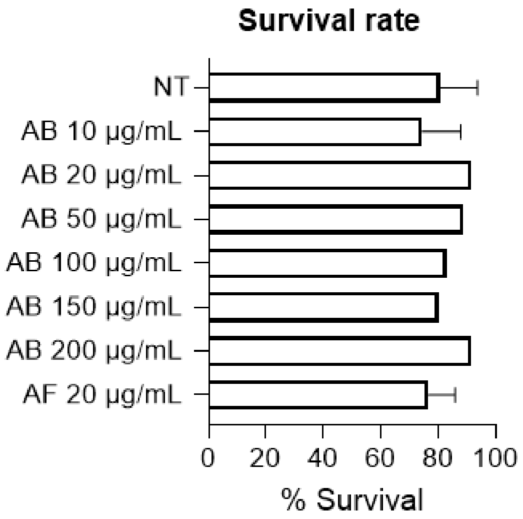

### Supplementary Tables

#### Table S1. Bacteria identified in the saliva of *Aedes aegypti.*

| Agar type | Bacteria | % of sequenced colonies | Accession numbers |
|---|---|---|---|
| **LB** | *Klebsiella* spp. | 31 | MK691689; KX214770; CP087580; MH680856 |
| | *Enterobacter* spp. | 16,9 | MK209686; OK326434; MN935016; MK764955 |
| | *Asaia* spp. | 16,9 | MK391152; MK128664 |
| | *Serratia marcescens* | 14,1 | KM091721; MH361385; CP055161 CP072199; MN788631; MT645673 |
| | *Serratia nematodiphila* | 12,7 | MT409573; |
| | *Microbacterium* spp. | 5,6 | MN922860; MT761105 |
| | *Chryseobacterium* sp. | 2,8 | MT539177 |
| **BHI** | *Enterobacter* spp. | 60 | OK326434; MN935016 |
| | *Klebsiella* spp. | 40 | MZ379584; KX214770 |
| **Blood** | *Enterobacter* spp. | 38,5 | MW435508; OK326434; MN935016 |
| | *Klebsiella* spp. | 38,5 | MK691689; KX214770 |
| | *Serratia marcescens* | 23 | CP055161; CP072199; MN788631 |

**Table S2. Bacteria identified in the saliva of *Culex pipiens.***

| Agar type | Bacteria | % of sequenced colonies | Accession numbers |
|---|---|---|---|
| **LB** | *Serratia marcescens* | 66,7 | KX904726; MH361385 |
| | *Serratia nematodiphila* | 33,3 | MT409573 |
| **BHI** | *Serratia marcescens* | 100 | KX904726; MT645673 |
| **Blood** | *Serratia marcescens* | 100 | CP072199; MN788631; MT645673 |

**Table S3. Sequencing input and output reads used for downstream analysis.**

| Samples | input | filtered | denoised | merged | output |
|---|---|---|---|---|---|
| Midgut | 94352 | 76181 | 76096 | 75660 | 75660 |
| Midgut | 116483 | 94833 | 94736 | 94098 | 94098 |
| Midgut | 108028 | 88778 | 88722 | 88221 | 88221 |
| Midgut | 133391 | 109364 | 109311 | 108237 | 108237 |
| Midgut | 153228 | 115421 | 115286 | 114807 | 114807 |
| Midgut | 114668 | 91291 | 91188 | 90347 | 90347 |
| NC | 123743 | 97161 | 96946 | 96550 | 96550 |
| Saliva | 115967 | 92701 | 92360 | 91185 | 91185 |
| Saliva | 153093 | 124181 | 123137 | 121554 | 121554 |
| Saliva | 135960 | 105277 | 104738 | 104164 | 104164 |
| Salivary Glands | 134445 | 115265 | 115041 | 114074 | 114074 |
| Salivary Glands | 128798 | 106331 | 106062 | 105183 | 105183 |
| Salivary Glands | 147122 | 115252 | 114567 | 113634 | 113634 |
